# Characterisation of the Avascular Mesenchyme during Digit Outgrowth

**DOI:** 10.1101/2025.03.17.643637

**Authors:** Cameron Batho-Samblas, Jonathan Smith, Lois Keavey, Noah Clancy, Lynn McTeir, Megan G. Davey

## Abstract

The avascular mesenchyme at the tip of the developing digit contributes to digit outgrowth and patterning, however, it has been poorly characterised. Using newly developed fate mapping approaches, tissue manipulation and single-cell mRNA sequencing data, we explore the transcriptional nature and developmental potential of this tissue. We find that the avascular mesenchyme is essential to normal segmental patterning of the digit and has a distinct transcriptional identity. In addition, we uncover an unexpected relationship between the unspecified tissue of the avascular mesenchyme and the committed phalanx forming region, which controls patterning, but not outgrowth of the digit. This multifaceted approach provides insights into the mechanics and genetic pathways that regulate digit outgrowth and patterning.

**Highlights:**

- The Avascular Mesenchyme of the developing digit contains a population of uncommitted cells, with a unique transcriptional identity.
- The AVM is critical to the normal outgrowth and patterning of digits.
- *HMGA1* expression is associated with the Avascular Mesenchyme

## Introduction

Digits- fingers and toes- are segmental structures made up along their length by a series of small bones called phalanges, separated by synovial joints. While the specification of digit number and identity are classic paradigms in developmental biology (Davey et al., 2018), the mechanisms which control the outgrowth and segmental patterning of digits are not clear. Altered digit anatomy is the most common form of congenital hand and foot birth defects, with important implications for welfare of both humans and animals (Stricker and Mundlos, 2011); brachydactyly, in which phalanges are small or malformed is estimated to occur in up to 2% of human live births (Temtamy and Aglan, 2008). Moreover, because our digits play such a central and important role in accessing and manipulating our environment, they are at huge risk of damage and their loss has profound functional consequences for daily life. In the US, it is estimated that a child receives emergency hospital care for a digit-tip injury caused by a door every 4 minutes; a third of these injuries will be digit amputations (Algaze et al., 2012). Furthermore, nearly half a million accidental finger amputations were recorded in US emergency departments between 1997-2016 (Reid et al., 2019). However, humans, like all other mammals, have a very limited ability to regenerate digits and as such digit regeneration has been intensely studied for many decades (Sensiate and Marques-Souza, 2019). The embryonic digit tip, in comparison has received little attention. We suggest that a study of the tissues and mechanics of the developing digit tip maybe informative in regenerative medicine.

The growing embryonic digit tip consists of five layers; the ectoderm, including the distal Apical Ectodermal Ridge (AER), a signalling centre which controls outgrowth of the digit through expression of Fibroblast Growth Factors (FGFs) (Sun et al., 2000), the Avascular Mesenchyme (AVM; also known as the non-condensed mesenchyme), the Marginal Sinus (MS), the Vascularised Mesenchyme (VM) and the Phalanx Forming Region (PFR; also known as the Digital Crescent) (Fig.1A,B). These layers interact dynamically, to drive concurrent elongation and segmentation of the digital ray along the distal-proximal axis, segmenting it into distinct chondrogenic elements that constitute the phalanges (Fig.1D, E). While the mechanisms that govern the specification of the digit primordia are highly conserved across vertebrate species, with most species exhibiting a pentadactyl limb structure (Saxena et al., 2017), the number of phalanges within each digit varies considerably. Cetaceans, for instance, display hyperphalangy, whereby each digit is elongated through the addition of extra phalangeal segments (Cooper et al., 2007; Richardson and Oelschläger, 2002). Additionally, In the chicken hindlimb the number of phalanges differs across digits; notably, digit IV consists of four phalanges and a terminal phalanx, making it a useful model in which to study digit outgrowth and patterning.

**Figure 1.**
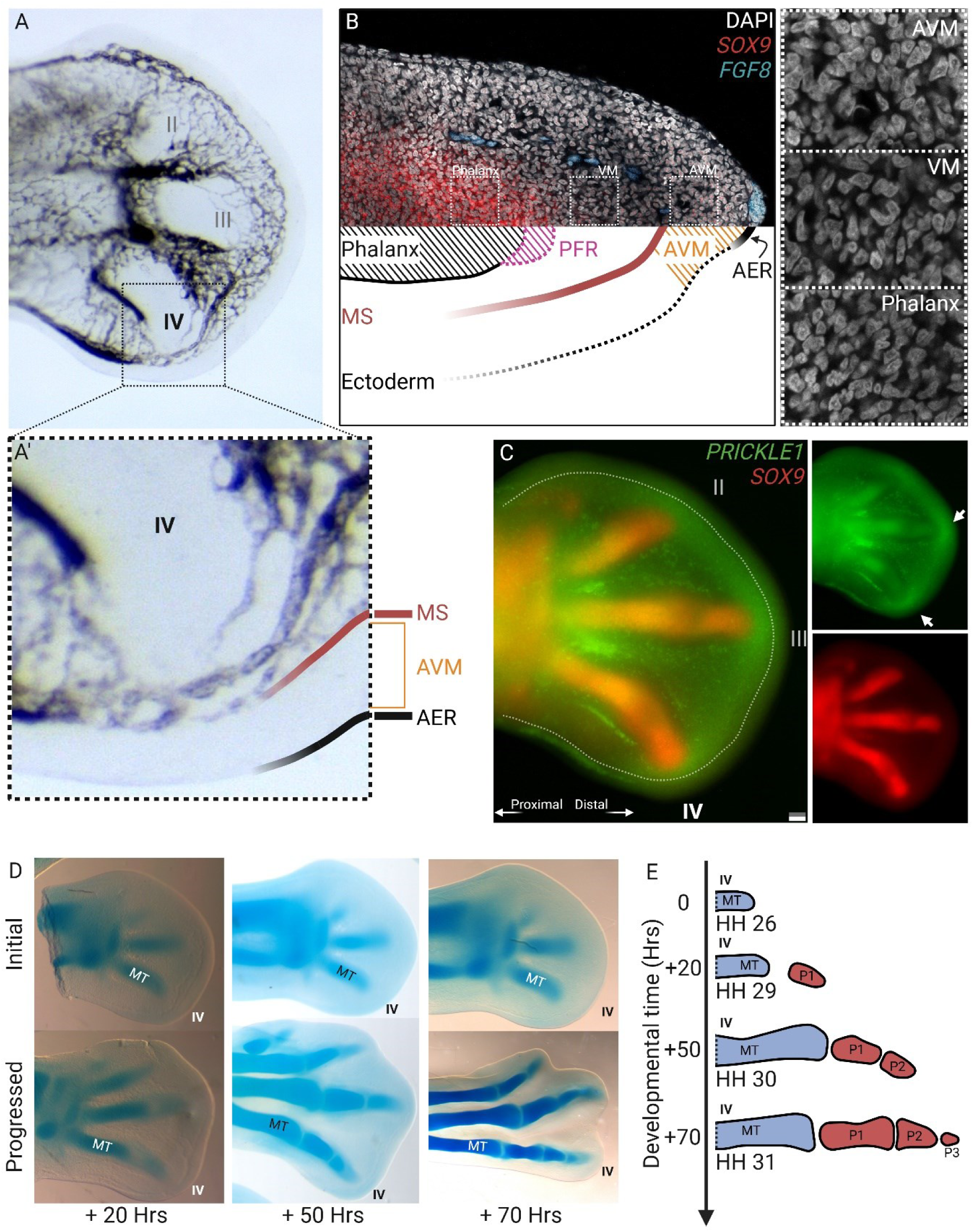
Characteristics of the avascular mesenchyme. (**A**) Vascular India ink injection of a HH 27 chick hindlimb autopod, with enlarged region within black dotted box (**A’**) (MS = Marginal Sinus, AVM = Avascular Mesenchyme, AER = Apical Ectodermal Ridge). (**B**) Top, fluorescent *in situ* hybridisation image of a section from a HH 27 hindlimb digit. Regions outlined by white dotted lines are enlarged to the right and only DAPI counterstain is shown (VM = Vascular mesenchyme). Bottom, schematic of digit tip anatomy along the proximal-distal axis (left-right) and dorsal-ventral axis (top-bottom). (**C**) Whole mount fluorescent *in situ* hybridisation of a HH 27 hindlimb autopod, showing the distally situated expression domain of *PRICKLE1*, which overlaps the AVM area, with strongest expression over the digits. White dotted line demarks the proximal boundary of the AVM. White arrows highlight regions of increased expression over the digit tips. (**D**) Cartilage staining on progressively older limbs (left to right), showing the approximate timings of phalanx formation in hours. (**E**) Schematic of phalanx formation timeline for digit (IV). MT = Metatarsal, P = Phalanx. Scale bars = 100 µm.

The PFR plays an essential translational role in interpreting signals from the interdigital mesenchyme into ‘digit identity’ which is primarily ascribed by the number of phalanges that form (Suzuki et al., 2008, Dahn and Fallon, 2000, Huang et al., 2016). The PFR has been thought to mediate the segmentation mechanism, thus, has been the focus of the majority of research on the digit tip (Huang et al., 2016, Witte et al., 2010, Montero et al., 2008, Hiscock et al., 2017). Importantly, the PFR is not the source of progenitor cells which form the digit, and, as seminal fate mapping experiments undertaken in chicken embryonic hindlimb have shown, the cells of the PFR are specified to form a specific phalanx, whereas the AVM is not (Suzuki et al., 2008). Cells of the PFR express *SOX9* (Fig.1B), an indication that cells in this domain are specified to form cartilage and tendon precursors. The AVM alone, however, can give rise to a digit, suggesting the AVM is the organiser of digit outgrowth and patterning, and contains a stem-cell like progenitor population (Suzuki et al., 2008). These fate maps have never been repeated, nor has the AVM been examined in further detail.

The AVM forms a peripheral layer approximately 100 ± 20 μm thick within the developing limb, enveloping underlying vascularised tissue (Caplan and Koutroupas, 1973). Although regions of avascularised mesenchyme are present in non-limb regions, its role in the limb exhibits distinct functional properties. At the distal end of the autopod the AVM forms a cap over the developing digits and interdigits, where it lies directly beneath the AER. Here the AVM displays a distinct gene expression profile, expressing a suite of genes that show spatial restriction to the distal mesenchyme, with genes such as *PRICKLE1*, *LHX2*, and *MSX1*. These genes have functional relevance to digit morphogenesis; for instance, *PRICKLE1* knockout mice form abnormal, fused, or shortened digits (Liu et al., 2014, Yang et al., 2013), highlighting the role of AVM-specific gene expression, and the AVM region itself, in digit formation.

Through the application of fate mapping, single-cell mRNA sequencing, and targeted structural manipulation, we demonstrate that AVM cells contribute not only to the formation of phalanges but also to synovial joints. Furthermore, we found that AVM function extends beyond progenitor cell contribution, playing a critical role in digit patterning, dependent on unknown signals from the proximal digit.

## Results

### The Avascular Mesenchyme is not determined, the Phalanx Forming Region is determined

First, we examined the expression pattern of *PRICKLE1*—a gene linked to digit outgrowth and patterning in mice (Liu et al., 2014; Yang et al., 2013)— in the autopod of an embryonic chick hindlimb. This revealed that *PRICKLE1* expression was not only present in the developing digital ray but was also specifically upregulated in the AVM overlying the PFR (Fig.1C). We then examined the timings of phalanx specification, building on earlier studies that identified Hamburger and Hamilton (HH) developmental stages associated with phalanx formation (Suzuki et al., 2008, Sanz-Ezquerro and Tickle, 2003). Our analysis revealed that in digit IV new phalanges are specified approximately every 20 hours, indicating a consistent segmentation period associated with the progression of distal outgrowth in the digit (Fig.1 D,E).

We then investigated the fate potentials of the cells from the AVM distal to the digital rays in the developing autopod. Through homotopic stage-matched grafting of the AVM and overlying AER distal to the digital ray of digit IV, we identified extensive contributions to the formation of both chondrogenic and connective tissue derived structures in the digits (n =6). This included the phalanges, synovial joints, and extra-digital mesenchyme, consisting of non-condensed mesenchyme surrounding the phalanges (Fig.2A-D). These observations are consistent with previous work by (Suzuki et al., 2008), which established that cells from the AVM migrate proximally to incorporate into the digit anlagen.

**Figure 2.**
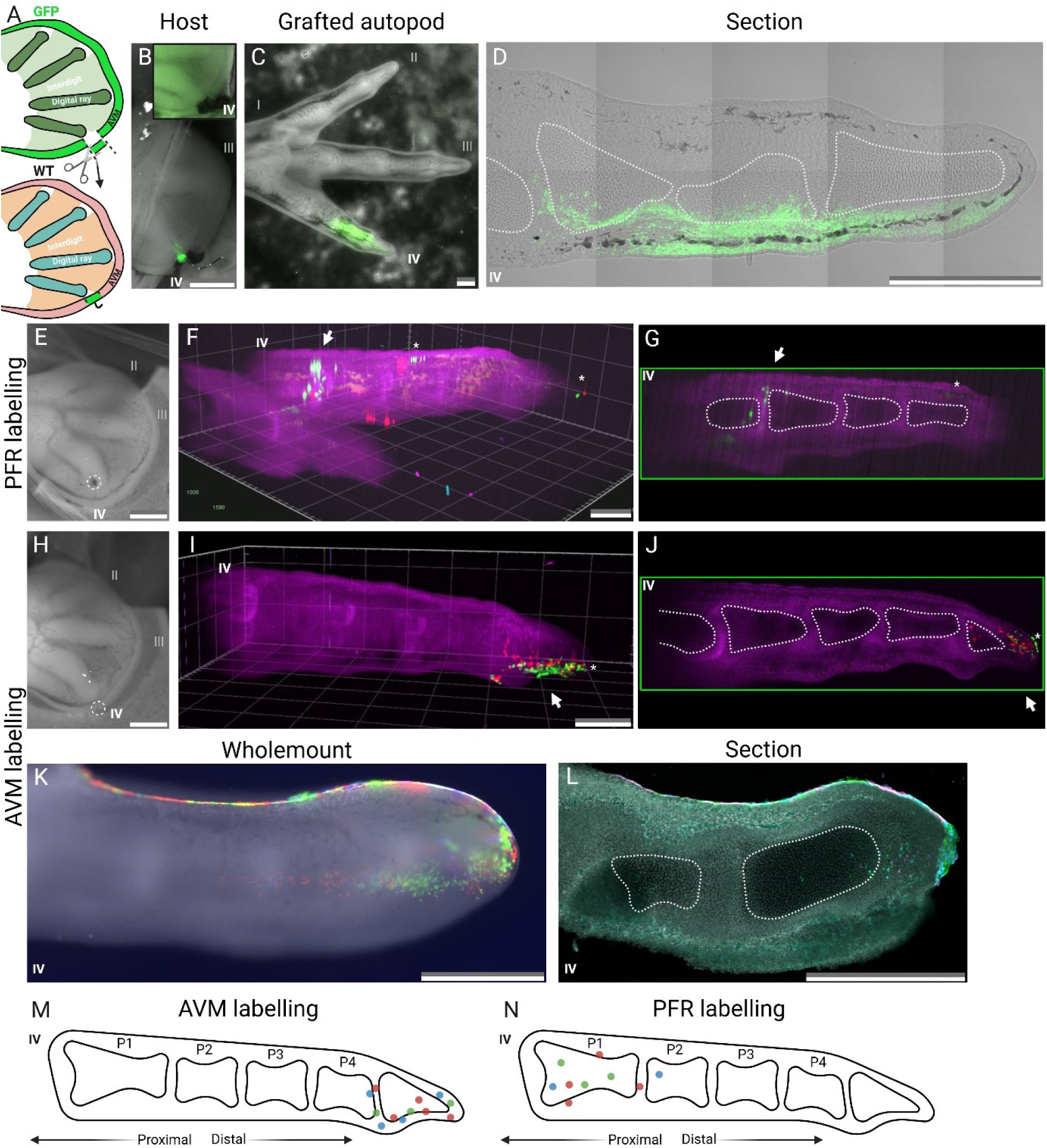
Fate potentials of the avascular mesenchyme and phalanx forming region. (**A**) Schematic showing a homotopic Avascular Mesenchyme (AVM) and overlying Apical Ectodermal Ridge (AER) graft, from digit IV in a HH 27 GFP transgenic embryo to a HH 27 WT host, as seen *in ovo* in (**B**). (**B**) Donor digit IV after dissection shown in black box (top right), host with donor tissue graft shown below. (**C**) The grafted cells (green) contribute to the terminal and penultimate phalanges in of an embryonic day (E) 11 digit IV. (**D**) Section of digit IV shown in **C**, confirming that the grafted cells are integrated into the digits and contribute to the phalanges, surrounding mesenchyme and the interphalangeal synovial joints whilst maintaining a population at the distal tip AVM. White dotted line demarks the phalangeal cells visible in plane. (**E, H**) Locations of beads soaked in TAT-Cre, labelling cells in either the phalanx forming region (PFR) (**E**) or AVM (**H**), are marked by white dotted circles. Beads were left at the indicated sites for 60 seconds in HH 27 hindlimb autopods before being removed. Lightsheet wholemount fluorescent image showing clones of labelled cells from the PFR (**F**) and AVM (**I**). Images to the right show optical sections through the middle of digit IV highlighting the predominant labelling of the proximal phalanx (P1) (**G)** and distal phalanges (P4 and terminal phalanx) (**J**). White arrows highlight labelled cells (red, blue, green) and asterisks denote ectodermal labelling. (**K**) Whole mount fluorescent image of clones from labelled AVM cells in E11 chick hindlimb digit IV and section of same digit (**L**). Schematic representations of the distributions of labelled clones resulting from labelling of the AVM (**M**) or PFR (**N**) highlighting the distinct domains of either region. Scale bars = 500 µm.

Predominately, cells from the grafted AVM tissue migrated proximally from the transplantation site, within the AVM, and into the digital tissues. Of note, a small subset of the grafted cells retained a distal position within the AVM, despite substantial distal-to-proximal outgrowth up to and including the latest timepoint of embryonic day (E11) (Fig.2C,D). This suggests distinct roles for the AVM during development; self-renewal and proliferation for digit-specific differentiation.

To investigate the developmental potentials of the AVM compared to the PFR we examined the difference in cell fates between these regions. This was achieved by sparse labelling of either the AVM or PFR at HH27, using the *Chameleon* transgenic chicken line. Through direct application of TAT-Cre delivered via Affi-Gel Blue beads (Oh et al., 2024), this approach enabled spatially and temporally specific permanent labelling of cells surrounding the beads (Fig.S1). By E11, following the formation of the terminal phalanx in all digits, labelled clones of cells originally within the PFR were almost exclusively localised within, and surrounding, the proximal phalanges P1 and P2 (n =8/8) (Fig.2E,G), excluding ectodermal labelling from initial penetration of the bead. Notably, labelling of the PFR revealed no evidence of a non-migratory subpopulation, as nearly all labelled cells had migrated into the proximal phalanges by E11, with no cells remaining within the PFR. In contrast, AVM-labelled cells produced results consistent with the homotopic grafting experiments (Fig.2B-D), whereby labelled cells were largely restricted to the terminal phalanx, phalanx 4 (P4) and phalanx 3 (P3), with a subset remaining in the AVM itself (Fig.2H-L; n = 6/6).

Additionally, optical sectioning and cryo-sectioning techniques confirmed that a subset of labelled cells from both the PFR and AVM became incorporated into the phalanges and synovial joints, rather than just within the surrounding mesenchyme (Fig2E-L;Fig.S2).

### Disruption to the Avascular Mesenchyme results in phalangeal patterning abnormalities

We demonstrated through homotopic grafting and labelling of the AVM that cells within the AVM contribute significantly to the tissues necessary for proper digit formation. Previous work has also shown that the AER, adjacent to the AVM, is essential for the distal-proximal outgrowth of the digits, through expression of diffusible factors, including FGF8, that repress differentiation and drive proliferation (Sun et al., 2000). However, surgical removal of the AVM and the overlying apical ectodermal ridge (AER) distal to digit IV at stage HH27 does not result in aborted outgrowth (failed development of any further phalanges or formation of a terminal phalanx).

At stage HH27, digit IV still exhibited a strong regenerative capacity, recovering outgrowth even after the loss of local AER signaling. Although outgrowth resumes, alterations to normal patterning were observed, affecting primarily the most distal phalanges (P3 and P4), while the proximal phalanges (P1 and P2) remained unaffected by the manipulation (n = 14/21, n = 19/21, respectively, Fig3A-H). These findings are consistent with previous homotopic grafting experiments and *Chameleon* fate mapping of the AVM. Of note, in multiple cases no skeletal patterning defects were observed in the manipulated digits (n = 6/21) and there was only a single case of no recovery of digit development.

The distal phalanges in manipulated digits were significantly elongated compared to their contralateral equivalents (p=0.0055), and in these cases digit IV lacked distal phalanges (Fig.3A-H). A minor phenotype also observed included the failure of interphalangeal joint 2 or 3 formation, leading to fusion of phalanges followed by the absence of P4 (n=3;Fig.3B-D). This was indicated by abnormal flaring within the phalanx shaft, a morphological feature typically restricted to the base and head of the phalanges. Fused digits displayed a significantly reduced head-to-shaft width ratio between 0.70 and 0.87, in contrast to unmanipulated control digits, which had ratios of 1.11 and 1.25 for P3 and P4, respectively (n=7; p=0.003, p=0.0006). This morphological feature aligns with the expected joint position when compared to the contralateral, unmanipulated side, suggesting that the phalanges maintain individual identify, but joint formation fails (Fig3D). In support of this hypothesis, RNA *in situ* hybridisation for the definitive interphalangeal joint marker *GDF5* (Storm and Kingsley, 1999) at E10 did not detect any joint progenitor cells within the shafts of the abnormal phalanges at locations along the proximal distal axis comparable to the contralateral digit (Fig3E-H), suggesting that joint progenitors were not specified.

**Figure 3.**
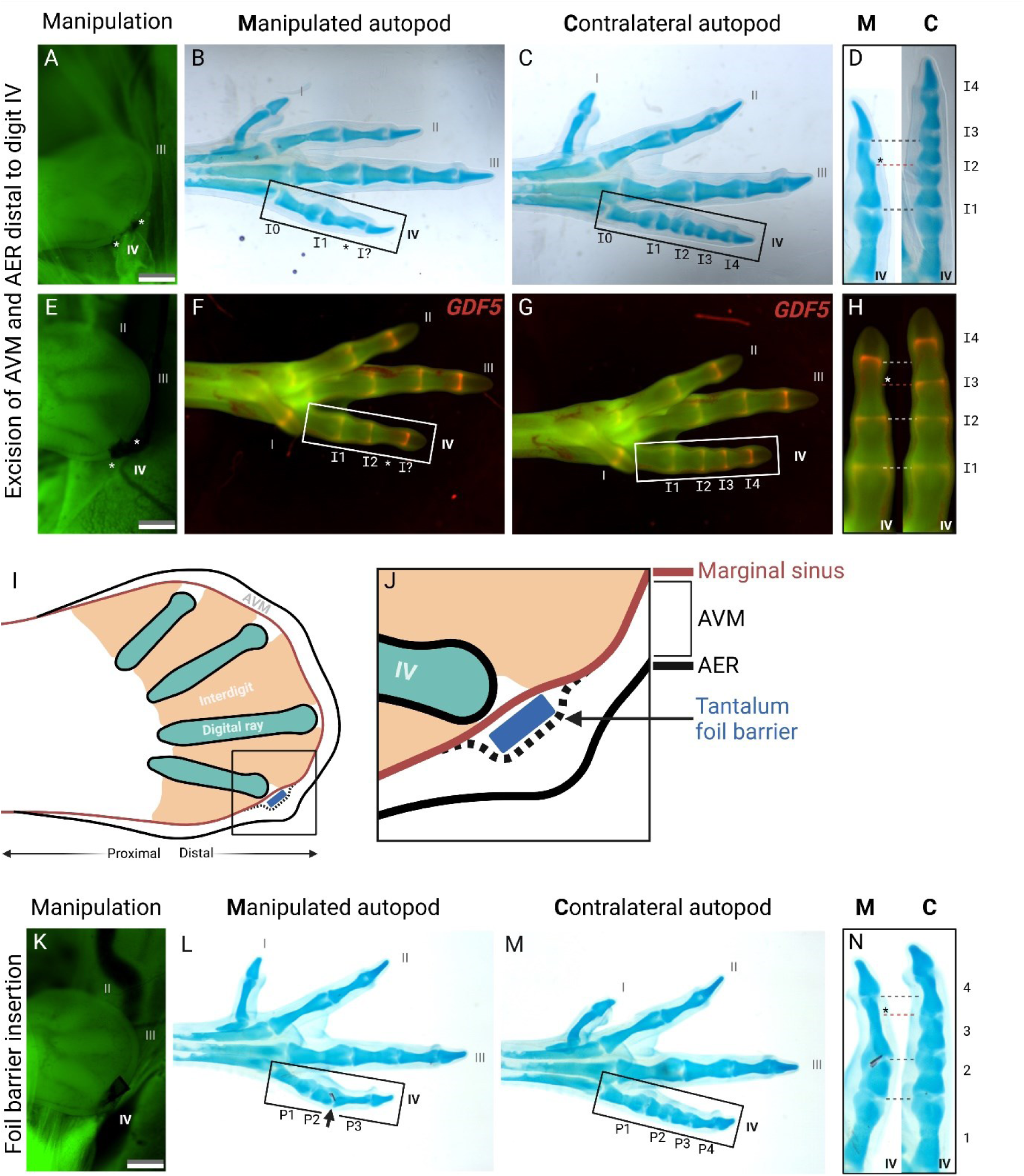
Disruption to the avascular mesenchyme results in abnormal joint patterning. (**A, E**) *In ovo* fluorescent images taken immediately post removal of the Avascular Mesenchyme (AVM) and overlying Apical Ectodermal Ridge (AER) in a HH 27 GFP chick hindlimb. White asterisks demark the breadth of tissue that was excised. (**B-C**) Cartilage staining of manipulated limb shown in **A** and control contralateral limb demonstrate removal of the AVM leads to elongation of the distal phalanx and missing penultimate phalanx (P4). (**D**) The morphology of the abnormal phalanx is suggestive of joint failure rather than elongation of the phalangeal element. (**F**, **G**) Wholemount fluorescent *in situ* hybridisation images of E11 chick hindlimb autopod, shown in **E**, demarking the definitive interphalangeal joint marker *GDF5* (Green = GFP, Red = *GDF5*). (**H**) Manipulated digit IV exhibited significant reduction in overall digit length (p = 0.007), elongation of P3 and absence of P4. White asterisk denotes expected position of interphalangeal joint 3 (I3) on the manipulated digit. (**I**) Schematic showing the location of the implanted tantalum foil barriers, as shown in **K**. Region in black box is enlarged on right (**J**), position of the foil barrier is at the proximal edge of the AVM, inserted into a slit made along the marginal sinus, distal to the digital ray of digit IV. Of note, the AER is undamaged. (**K**) *In ovo* fluorescent image of HH 27 GFP chick hindlimb, representative of the tantalum foil barrier insertions. (**L-N**) The same embryo sacrificed at E11, cartilage stained and cleared for imaging. The manipulated side contains the original tantalum foil insert (demarked by black arrow) within interphalangeal joint 2. Results in elongation of the phalanx immediately distal to the foil barrier (P3) and loss of P4, when compared to the contralateral control. Scale bars = 500 µm.

Of note, the manipulated digits which recovered all formed terminal phalanges (n=20/20), which suggests that the endogenous patterning program was successfully re-established after the manipulation. Additionally, unlike the interphalangeal joints, the terminal phalanx did not exhibit delayed development and appeared morphologically normal when compared to the contralateral digit.

In summary, post recovery the digits exhibited two primary phenotypes: a reduction in the number of phalanges by one or two elements, both of which were accompanied by a significant reduction in length of the manipulated digit (p=0.007). These phenotypes are unlikely to represent homeotic transformations of digit identity as in some cases the remaining distal phalanges displayed signs of joint formation failure, and subsequent fusion of phalanges rather than remodelling and elongation of phalanx morphology. While the overall phalanx morphology was largely normal, the persistent failure to separate into distinct phalangeal elements suggests a disruption in the segmentation process following the removal of the AVM and AER.

We next examined signalling between proximal tissues and the AVM, as while previous studies have demonstrated that the interdigit region proximal to the PFR signals to the PFR in order to determine digit identity (Dahn and Fallon, 2000; Huang et al., 2016), signalling from proximal tissue to the AVM remains unexplored. To test this we inserted tantalum foil barriers at the proximal boundary of the AVM, adjacent to the marginal sinus, while maintaining an intact AER. The barriers were cut to an average width of 164.25 µm and were positioned directly distal to the developing digit IV in stage HH 27 embryos (Fig.3I-K). Successful insertion of the foil resulted in either a total loss of phalanges distal to the foil, but development of a terminal phalanx (n = 2/12), or reduction in number of phalanges distal to the foil. Additionally, phalanges distal to the foil showed a significant increase in length (n = 8/12) (p=0.005) (Fig.3K-M). In digits in which the foil was displaced, fell out, or limbs in which only a slit was made but no foil placed, digit growth and phalanx morphology was normal (n=13; not shown). Likewise, foil placements that deviated by small margins latterly from the centre of the developing digit did not result in alterations to patterning. In such cases the foil barrier was found to be located adjacent to the phalanges (in the non-condensed mesenchyme; n=3). This was in contrast to those placed at the midline, whereby the foil was incorporated into the digits, specifically located at, or immediately adjacent to, the 3^rd^ interphalangeal joint (n=8/21).

Despite the alteration in patterning caused by the foil, the formation of a terminal phalanx was observed in most cases (n=11/12). The presence of a terminal phalanx suggests that the manipulated digit still terminated patterning normally, in a non-abortive fashion. Additionally, the proximal phalanges (P1 and P2) were unaffected by the insertion of the foil barrier (n=10/12). This is consistent with the findings from homotopic grafting AVM experiments and AVM fate mapping using the *Chameleon* transgenic line. This suggests that the disruption in joint formation may be caused by of the manipulation of the AVM cells and supports the conclusion that proximal tissue may interacts with the AVM for phalangeal/joint patterning.

### Ectodermal signalling to the Avascular Mesenchyme is essential for proper phalangeal patterning

Ectodermal synthesis of hyaluronic acid (HA) has been implicated in the formation of avascular zones across multiple embryonic tissues, (Sobolewski et al., 2005; Mcclean and Rogers, 1945; Lin et al., 2022), including in the developing limb bud epithelia, where high HA levels are sufficient to induce de novo avascular zones in the limb bud (Feinberg and Beebe, 1983). Recently it has been shown that regeneration of amputated mouse digit tips requires HA and can initiate the rescue of non-regenerative amputations (Mui et al., 2024). To investigate the role of avascularity in the AVM with respect to digit morphogenesis, we sought to promote vascularisation within the AVM by the inhibition of local HA synthesis. This was achieved by the insertion of 4-methylumbelliferone (4-MU) beads into the AVM immediately distal to the digital ray of digit IV at HH27 (Fig.4A). Analysis of the AVM 6 hours post implantation of 4-MU bead revealed a significant reduction in the width of the AVM distal to the manipulated digit, suggesting that ectodermal HA expression plays a role in maintaining the AVM (Fig. S3).

**Figure 4.**
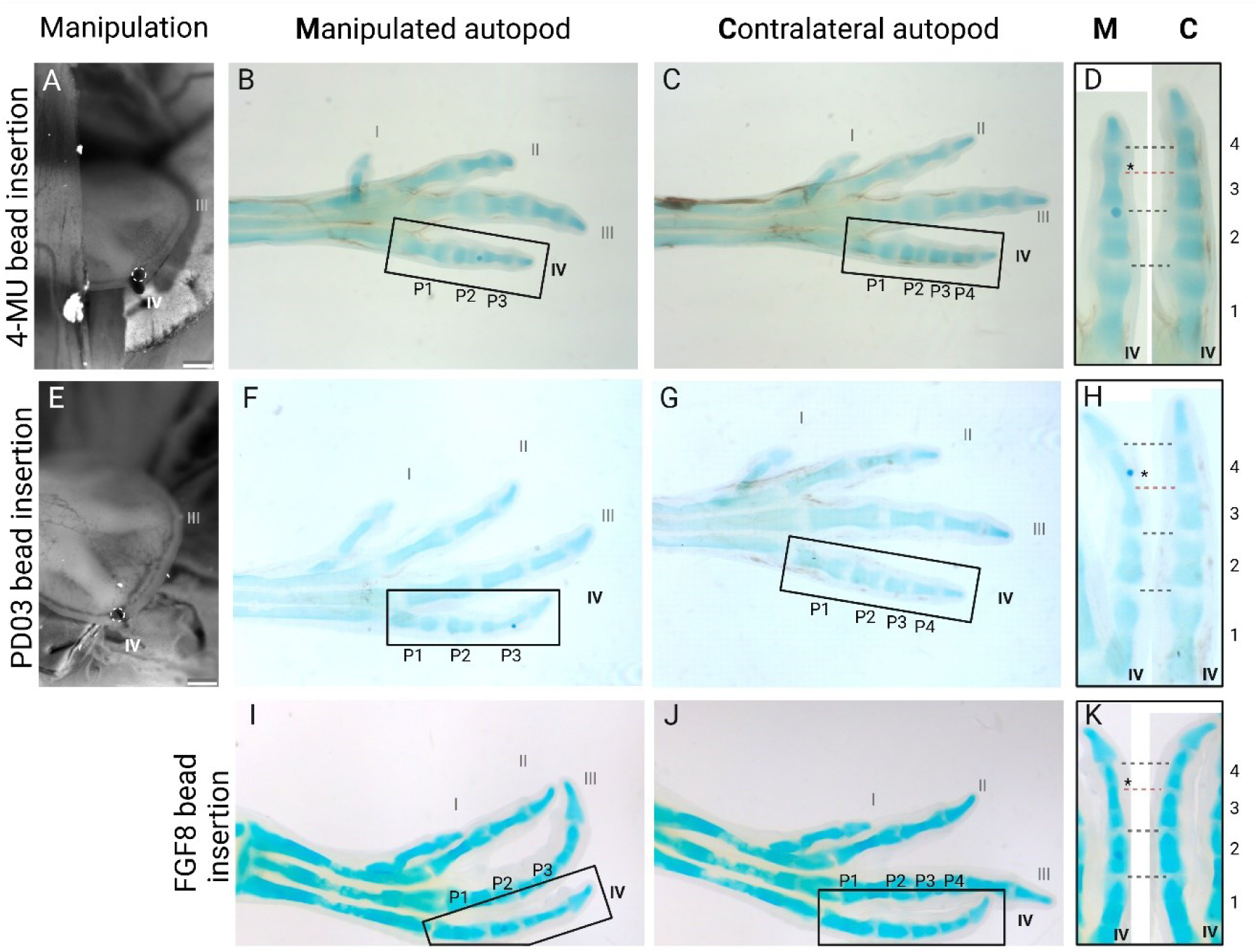
Disruption of ectodermal signalling to the avascular mesenchyme results in abnormal phalangeal patterning. (**A**) Implantation of a bead soaked in 4-MU into the AVM distal to digit IV in a HH 27 embryo. White dotted circle denotes location of bead. (**B-D**) The embryo shown in **A** at embryonic day (E) 11 after cartilage staining and clearing. Black asterisk denotes expected position of interphalangeal joint 3 on the manipulated digit. The manipulated digit IV does not have phalanx 4 (P4) and exhibited elongation of P3. (**E**) Implantation of a bead soaked in a pharmacological inhibitor of MEK/ERK signalling, PD0325901, into the AVM distal to digit IV in a HH 27 embryo. (**F-H**) At E11 fusion between P3 and P4 was observed in the manipulated side when compared to the contralateral. (**I-K**) Implantation of a bead soaked in FGF8 into the distal mesenchyme posterior to digit IV in a HH 27 embryo was performed which results in elongation of P3 and absence of P4 when compared to the contralateral. Scale bars = 500 µm.

4-MU treatment produced skeletal developmental defects observable at E11. Specifically, manipulated digits exhibited altered digit patterning, with an absent distal phalanx (P4) and an elongated P3 (n=3/4; p=0.029; Fig.4, B-D). Furthermore, the overall length of digit IV was significantly reduced by an average of 8.3% compared to the unmanipulated contralateral digit (p=0.020). Despite these defects, a terminal phalanx still formed. These findings suggest that 4-MU may result in impaired cellular proliferation, reduced migratory capacity or a reduction in BMP signalling within the AVM, contributing to decreased digit outgrowth.

The other major signal from the overlying ectoderm, specifically the AER, is FGF8, which signals through MEK/ERK. The MEK/ERK signaling pathway plays a critical role in digit morphogenesis, integrating inputs from several major developmental pathways, including EGF and Neurotrophin (Ullrich and Schlessinger, 1990; Seger and Krebs, 1995; Kaplan and Miller, 2000). However, FGF8 represents the primary signaling factor that signals through MEK/ERK with respect to the distal digit. To examine the role of MEK/ERK signaling within the AVM, we locally inhibited this pathway using the potent MEK inhibitor PD0325901. The beads were positioned distal to digit IV at stage HH27, which by E11 resulted in disrupted distal phalangeal patterning, particularly affecting joint formation in the distal phalanges (P3 and P4; n = 4/10; Fig.4E-H). In these cases, the heads and tails of the distal phalanges displayed morphological abnormalities and lacked distinct interzones, leading to fusion of distal phalangeal elements, including the terminal phalanx (n=2/10). This resulted in significant elongation of P3 (p=0.005) and, notably, the affected digits were significantly smaller than their contralateral counterparts (p=0.019) suggesting a phenotype consistent with reduced FGF8 signaling. The results of PD0325901 bead implantation indicate that while phalangeal segmentation initiates it does not fully complete.

Interestingly, beads soaked in FGF8 and implanted into the distal mesenchyme posterior to digit IV produced a phenotype similar to that observed with MEK/ERK pathway inhibition. This phenotype included elongation of the phalanx distal to the bead, absence of P4, and formation of a terminal phalanx (n = 4/6; Fig.4I-K). Despite FGF8’s established role in promoting cell proliferation and digit outgrowth (Sun et al., 2000), the manipulated digits were significantly shorter than their contralateral counterparts when compared to unmanipulated controls (p=0.005), closely mirroring the effects seen with PD0325901 treatment. These findings suggest that a balance of FGF8 levels, rather than exceeding a threshold, is essential for normal digit morphogenesis.

Of note, the rate and extent of diffusion for each of the pharmacological agents were not quantified, making it not possible to determine whether adjacent digit regions were affected or contributed to the observed phenotypes. However, minor perturbations to bead placements resulted in a lack of gross phenotypic changes which suggests that only a small region surrounding the bead was affected.

### Transcriptional profiling of the Avascular Mesenchyme

To investigate the transcriptional profile of the AVM during digit development, we analysed single-cell mRNA sequencing data from chick hindlimb autopods at 25HH and 31HH (Feregrino et al., 2019). Post quality control filtering, the combined datasets consisted of 12,207 cells. This was followed by unbiased cell clustering, which identified many of the expected cell types from the chick autopod including epithelial cells (Cluster 4), chondrocytes (Cluster 1), and interdigital mesenchyme (Clusters 6 and 8) (Supplemental Information 2, Supplemental Table 1) (Fig.5A, B). We focused the analysis of the sequencing data to identify a population of cells enriched in the digit-specific stemness factor *PRMT5. PRMT5* is a crucial factor in the maintenance of the digit progenitor population, where in mice its loss of function results in significant reductions in digit outgrowth as well as patterning defects (Norrie et al. 2016). We and others (Suzuki et al. 2008) have demonstrated through fate mapping experiments that the progenitor population to the digits is within the AVM, thus, given *PRMT5’s* spatial and functional specificity to the AVM it was used to screen the mesenchymal clusters.

**Figure 5.**
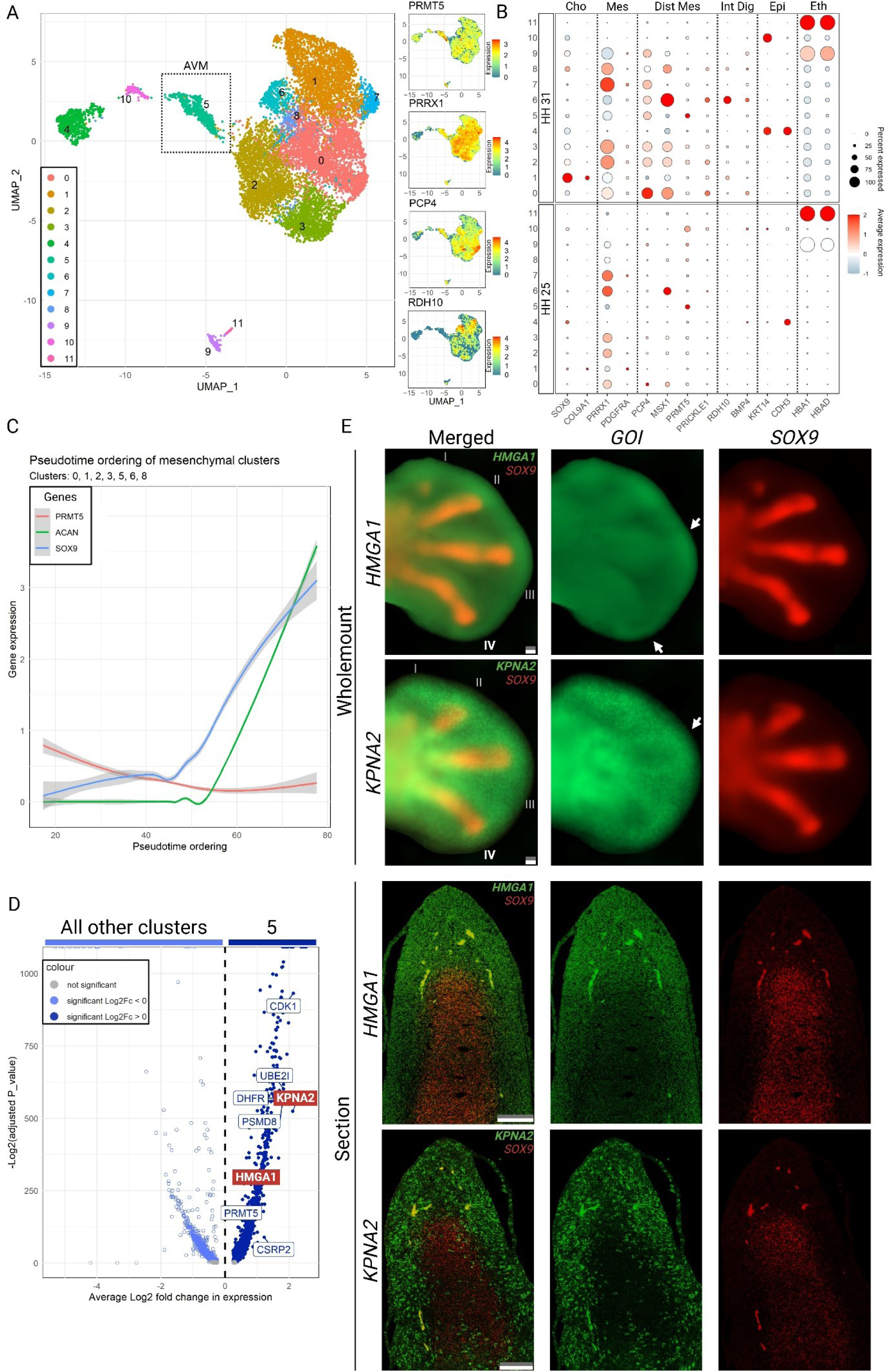
Characterisation of the transcriptional profile of the avascular mesenchyme. (**A**) UMAP representation of the combined HH 25 and HH 31 datasets (Feregrino et al. 2019) totalling 6,769 and 5,438 cells respectively. Colouring of the cells indicate the result of unsupervised clustering methods. To the right are feature plots showing the expression patterns for marker gene expression on the UMAP embedding for the Avascular Mesenchyme (AVM), mesenchyme, distal mesenchyme and interdigit (top to bottom). (**B**) Dotplot of known marker genes for the broad limb populations identified within the mRNA single cell sequencing data (Cho = Chondrocyte, Mes = Mesenchyme, Dist Mes = Distal Mesenchyme, Int Dig = Interdigit, Epi = Epithelium and Eth = Erythrocyte). All clusters were annotated with the exception of cluster 10. (**C**) Trajectory analysis on the mesenchymal clusters generated a pseudotime ordering to the cells, initiating in cluster 5 (AVM) to cluster 0 (distal mesenchyme) and terminating in cluster 1 (chondrocytes). Gene expression was plotted over this ordering recapitulating gene expression patterns of cells migrating from the AVM. (**D**) Volcano plot showing genes significantly enriched and upregulated for cluster 5, identified through differential expression analysis. (**E**) Fluorescent *in situ* hybridisation of genes highlighted in red in **D**. Both wholemounts and sections were of approximately HH 26-27 (E 6) hindlimbs, where *HMGA1* showed the greatest enrichment to the AVM. White arrows highlight regions of increased expression over the digit tips. *GOI* = gene of interest Scale bars = 100 µm.

Differential expression analysis identified Cluster 5 to be significantly enriched in *PRMT5* expression (Fig5B), exhibiting a 1.8x fold increase in *PRMT5* when compared to all other clusters (p=0.00), a trend consistent across both 25HH and 31HH datasets. Additionally, Cluster 5 expressed the distal mesenchyme markers *PCP4* and *LHX2*, however, at significantly lower levels than within the identified distal mesenchyme cluster (Cluster 0; Fig.5B) (Supplemental Information 2). Furthermore, Cluster 5 did not show significant association with many other prominent distal mesenchymal markers including *MSX1* and *PRICKLE1*.

As earlier experiments have demonstrated (Fig.2), the AVM has the developmental potential to give rise to many of the digit structures, including the chondrocytes that form the digital rays. This is reflected in the transcriptomic data where trajectory analysis performed on a mesenchymal subset of the data identified a developmental trajectory originating in Cluster 5 and terminating in the chondrocyte population, Cluster 1 (Fig.5C). The pseudotime ordering correctly recapitulated the progressive loss of the distal AVM marker *PRMT5*, and showed the increasing expression levels of *SOX9* as the cells progress towards a chondrogenic fate. This was acquired before *ACAN*, consistent with the known fate of the AVM cells (Lefebvre and Smits, 2005; Bi et al., 1999).

Differential expression analysis on Cluster 5 identified 693 significantly upregulated genes (fold change >1.5 and adjusted p-value <0.05) when compared to other clusters in the combined dataset. The top 20 most enriched and upregulated genes within Cluster 5 were predominately known to be involved cell cycle functions as positive regulators of proliferation, including *DHFR*, *CDK1* and *DSCC1* (Zhu et al., 2023; Gangjee et al., 2007; Nurse, 1990; Bermudez et al., 2003), and most did not have any reported functional relevance to limb development. This was reflected in gene set enrichment analysis. When using the top 200 positively differentially expressed genes the only enriched biological process term was translational initiation (Fig.S5).

Although no gene expression was found to be entirely unique to Cluster 5, several genes - *KPNA2*, *HMGA1*, and *CSRP2* - exhibited elevated expression and specificity to Cluster 5 within the combined dataset (Figure 5, D; Figure S7). *HMAG1* and *KPNA2* were validated by fluorescent RNA *in situ* hybridisation chain reaction (HCR), enabling visualisation of expression domains in 27HH chick hindlimb autopods (Fig.5E, Fig.S7, S8). The identified genes showed prominent expression in the developing autopod, however, *HMGA1* exhibited the most restricted expression to the distal mesenchyme.

The *HMGA1* expression domain, approximately 300 µm in width, spanned the distal perimeter of the autopod, covering both the AVM and vascular mesenchyme regions while displaying reduced presence in the digital rays. Notably, this expression pattern showed high levels of similarity to *PRICKLE1*, a gene with functional relevance in digit development, however, this was not reflected in the single-cell mRNA sequencing data. Cluster 5 did not exhibit high levels of expression of *PRICKLE1*, which may indicate cellular heterogeneity within AVM and distal mesenchyme. Of note, like *PRICKLE1*, *HMAG1* also displayed periodic increased levels of expression across the autopod, concurrent with the digits, suggestive of an active role in digit morphogenesis (Fig. S10).

Taken together, Cluster 5 likely represents a heterogeneous grouping of cells. Cluster 5 likely includes the AVM, indicated by its comparatively high levels of *PRMT5* expression and the distal expression patterns of *HMGA1* and *KPNA2*. However, the lack of significant expression of other distal mesenchyme markers suggest that Cluster 5 was not wholly constituted by transcripts from the AVM. Further experimental investigation is required to resolve this cellular heterogeneity.

Additionally, *HMGA1* has been identified as a potential stem cell maintenance factor in the chick neural tube, and has confirmed roles supporting initiation of migration and subsequent separation from the progenitor pool (Gandhi et al., 2020). This raises the possibility of a similar role for *HMGA1* in chick digit development, particularly in the function of the AVM progenitor pool.

## Discussion

Here we provide new insight into the role of the AVM in digit morphogenesis, offering evidence for its contributions to both digit tissues and in joint specification within developing digits. Fate mapping experiments revealed that cells within the AVM contribute to essential digit tissues, notably including both the phalanges and the interphalangeal synovial joints—elements critical to the proximal-distal patterning of digits. Additionally, a subset of cells within the AVM remained spatially restricted to the AVM over the course of digit morphogenesis. This is a behaviour which suggests an AVM-specific program of self-renewal, reinforcing its role as a proliferative digit progenitor population. The findings from this study support the view that the AVM’s unique properties underlie its developmental role, operating separately from, but in a coordinated fashion with, the PFR.

AVM removal experiments further underscored the AVM’s significance. Intriguingly the resulting phenotypes do not replicate the outcomes of AER removals, however, this is likely due to the larger wound size and subsequent activation of a wound response enabling effective regeneration. Variability in the number of phalanges formed post-AVM removal may be attributed to differences in developmental timing between embryos or differences in the extent of tissue excision. Of note, the elongated phalanx distal to the manipulation closely resembled those observed in foil barrier experiments, which suggests a common underlying mechanism: disruption of proximal-distal signaling to the AVM. In both cases the signalling environment of the AVM is not accessible to the proximal digit, including the PFR, and from these experiments it is apparent that the temporary breaking of signalling leads to permanent change in the behaviour of the AVM. In both cases the PFR is undamaged, and yet once the digit has grown past the foil barrier, or the AVM has regenerated, normal patterning does not resume.

Our findings align with previous work by (Suzuki et al., 2008), which demonstrated that AVM grafts from digit III to the trunk of embryonic chick results in a phenotype that matches the AVM removal and barrier insertion phenotypes: formation of a single phalanx and a terminal phalanx. This represents the intrinsic and autonomous patterning programmes within the AVM when receiving no external (proximal) signals; a digit without identity. This opens up the possibility that the AVM removals/barrier insertions result in a loss of proximal signalling cues to the AVM, leading to loss of digit identity. As the PFR has remained unmanipulated this suggests that the AVM and PFR must communicate during normal digit morphogenesis to confer identity.

These findings coupled with the observed joint fusion in manipulated digits suggests that the AVM likely influences joint formation by mediating the specification of joint progenitor cells. Even with an intact PFR and a regenerated AVM joint specification was compromised, pointing to an essential ongoing role for the AVM in regulating this process. Additionally, pharmacological manipulation with 4-MU, which we show to disrupt the avascular state of the AVM, exhibited instances of joint failure, again indicating the AVM’s role in specifying joint progenitors. This finding points to the avascular, hypoxic environment within the AVM as potentially key to its function, possibly influencing region-specific gene expression patterns necessary for joint progenitor development.

Our investigation into the role of FGF8, a key signaling molecule from the AER, on AVM function revealed that both inhibition (using PD0325901) and overexpression of FGF8 led to similar phenotypic outcomes, a similar finding to Sanz-Esquerro and Tickle 2003. This unexpected lack of a binary response suggests that FGF8 may function within an oscillatory signaling network or mediate cell migration. Perturbations to this network, whether through increased or decreased FGF8 levels, could disrupt the timing and progression of this oscillatory mechanism or the direction of movement of cells within a morphogenic field, ultimately resulting in similar phenotypic consequences.

Additionally, our bioinformatics analysis identified a novel marker of the AVM and distal mesenchyme, *HMGA1,* which was enriched in and significantly upregulated within Cluster 5. Although *HMGA1* expression overlaps with known distal markers like *MSX1* and *PRICKLE1*, Cluster 5 cells did not universally co-express these markers, indicating that the distal mesenchyme, including the AVM, may be more heterogeneous than previously recognised. These findings imply that the AVM comprises subpopulations with potentially distinct roles in digit morphogenesis, and that the distal mesenchyme may contain more complexity than what can be observed morphologically.

Taken together, our findings emphasise the AVM’s contributions to digit outgrowth and joint patterning/specification and will be a useful addition towards modelling digit outgrowth and patterning undertaken by other (Scoones and Hiscock 2020; Grall et al. 2023). The AVM appears to orchestrate a range of developmental processes beyond just self-renewal and proliferation, suggesting a dynamic interplay between signaling gradients, hypoxic niches, and intrinsic cellular programs. The role of HA in development appears to parallel the role of HA in regeneration and may play a role controlling the interplay between tissue stiffness (Mui et al., 2024) and cellular flow (Parada et al., 2022) during digit outgrowth and morphogenesis. These insights deepen our understanding of digit morphogenesis and underscore the AVM’s role as a complex component, tightly integrated into the functions of the developing limb tissues.

## Materials and methods

### Chicken husbandry

All animal management and maintenance were carried out under UK Home Office license (PP8571233- Breeding and Maintenance of GA poultry lines) and regulations. Experimental protocols and studies were approved by the Roslin Institute Animal Welfare and Ethical Review Board Committee. Fertilised eggs from the ISA Brown, Roslin Green (Cytoplasmic eGFP), Flamingo (TdTomato) and Chameleon (Cytbow) lines were incubated at 38°C until they had reached the desired developmental HH stage. The eggs were windowed and prepared for manipulation as per Tiecke and Tickle 2007. Once manipulated they were sealed with tape and re-incubated at 38°C until the desired HH stage. Embryos were culled by a Schedule 1 culling.

### Affi-gel bead implantation

20 µl of Affi-Gel Blue Gel beads (BioRad, 1537302) were suspended in phosphate buffered saline (PBS) and stored at 4°C. Prior to use, the beads were transferred to a humidified plate as per Tiecke and Tickle (Tiecke and Tickle, 2007) and soaked in TAT-Cre Recombinase (EMD Millipore, SCR508, 1500 units), PD0325901 diluted in DMSO (at concentrations of 5 mM or 1 mM, 100% and 10% DMSO respectively), FGF8 (Gibco, PHG0274) (1 µm/1 µl), or 4-MU (Stratech, B6001) diluted in DMSO (at concentrations of 1 mM or 0.1 mM and 10% and 1% DMSO respectively). The beads were soaked in the respective reagent for a minimum of 1 hour at 4°C. A slit was made following the marginal sinus at the desired site using a tungsten dissection needle, creating a loop of tissue. The beads were then picked up by penetrating them with an electrolytically sharpened tungsten dissection needle, briefly air-dried to allow shrinkage, and inserted into the loop of tissue, where they were left in place unless otherwise specified. The eggs were then resealed with tape and incubated at 38°C in a humidified, light-free environment until they reached the desired Hamburger and Hamilton stage (Hamburger and Hamilton 1951).

### AG1-X2 bead implantation

4-MU was prepared by diluting in DMSO to concentrations of 1 or 0.1 Mm. AG1-X2 anion exchange beads (Bio-Rad) were soaked in the 4-MU solution for a minimum of 1 hour at room temperature and under light-free conditions. Following incubation, the beads were washed twice in media and subsequently kept in media at room temperature. At the appropriate embryonic stage, a slit was cut at the target site for bead insertion using a fine tungsten needle. A single bead was then transferred using a P20 pipette, placed near the slit, and inserted into the site. The eggs were then resealed with tape, and the embryos were incubated at 38°C until they reached the desired Hamburger and Hamilton stage (Hamburger and Hamilton 1951).

### Grafts and removals

Stage-matched homotopic transplantations were performed using Roslin Green, Flamingo, and wild-type ISA Brown chick embryos. Host sites were dissected and removed using a fine tungsten dissection needle, and equivalent donor regions were carefully dissected and positioned into the host site using filter paper (Merk, HABP04700). The donor tissues were secured in place with 0.02 mm oxidized nickel-chrome wire, ensuring that the apical ectodermal ridge (AER) was oriented away from the host site to match its endogenous orientation.

The eggs, both donors and hosts, were then resealed with tape and incubated at 38°C in a humidified, light-free environment until they reached the desired Hamburger and Hamilton stage (Hamburger and Hamilton 1951).

### Tantalum foil insertion

Fine strips of tantalum foil (0.025 mm thick) were cut to an average width of 164.25 µm using a scalpel. A slit was made following the marginal sinus at the desired site using a tungsten dissection needle. The Tantalum foil strip was inserted into the slit, ensuring it passed through both the dorsal and ventral faces of the digit.

### Vascular ink injection

Filtered India ink was diluted at a 1:5 ratio in PBS. Chick embryos at the appropriate Hamburger and Hamilton developmental stage were windowed to allow access to the extraembryonic vasculature. India ink was aspirated into a pulled glass capillary needle and carefully injected into a chorioallantoic membrane vessel by mouth-pipette. Once the ink had fully perfused the vasculature of the embryo, they were quickly dissected into ice-cold DEPC PBS.

Embryos were fixed overnight in 5% trichloroacetic acid (TCA), followed by three washes in 70% ethanol for 10 minutes each with gentle rocking. After washing, embryos were incubated in 100% ethanol overnight and subsequently cleared in methyl salicylate (Merk, M6752) for imaging and further analysis.

### Alcian green cartilage staining

Dissected embryos were fixed overnight in 5% TCA at room temperature, whilst rocking. After fixation, the embryos were washed twice in 70% ethanol for 5 minutes while rocking, followed by a 1 x 5 minute wash in 70% EtOH (Ethanol): 1% hydrochloric acid (HCL). The embryos were incubated overnight in 70% EtOH: 1% Alcian green: 1% HCL. Embryos were washed for 3 hours in 70% EtOH: 1% HCL followed by a quick wash in 70% EtOH. To complete the process, embryos were gradually dehydrated to 100% EtOH and cleared in methyl salicylate for imaging.

### Sectioning

Limbs were dissected into DEPC PBS where they were then fixed in 4% PFA for 1-2 hours at room temperature. Following fixation, the samples underwent 2 x 5 minute washes in PBS. The samples were washed in a 5% sucrose:PBS solution for 15 minutes whilst both rocking and on ice. Samples were then washed in 20% sucrose:PBS solution for 25 minutes, rocking and on ice. Samples were immersed in a sucrose-gelatin (Sigma-Aldrich, G2500) solution and incubated at 37°C for 30-40 minutes. Following this, the samples were placed in a petri dish filled with the sucrose-gelatin solution, covered, and kept at 4°C for 30-40 minutes. The samples were then cut out of the sucrose-gelatin block and mounted to a thin section of cork using Optimal Cutting Temperature compound (OCT). The mounted samples were frozen in −80°C isopentane and stored at −80°C until sectioning.

The specimen and chamber temperatures were pre-cooled to −25°C, as were the chuck and cutting blade. Samples were affixed to the chuck using OCT on the cork side. A cut depth of 10 μm was used for sectioning. The resulting sections were placed on Superfrost slides and left covered at room temperature for 2-3 hours. Subsequently, the excess sucrose-gelatin was removed from the slides through a wash PBS at 37°C PBS (water bath) for 30 minutes.

The sections were mounted using antifade Prolong Gold (Thermo Fisher, P36930) and a number 1 cover slip (22 mm x 50 mm) and then stored in the dark at room temperature. Images were captured using a Zeiss LSM 880 confocal microscope.

### Hybridisation chain reaction RNA *in situ* hybridisation

Limbs were dissected in Diethyl pyrocarbonate-treated phosphate-buffered saline (DEPC PBS) and transferred for fixation into 4% Paraformaldehyde (PFA) for a duration of 1-2 hours at room temperature. The samples were then washed in PBS for 2 x 5 minute washes before being dehydrated through sequential 5-minute washes in MeOH (Methanol):PBS solutions with concentrations of 25%, 50%, 75%, and 100% whilst rocking and on ice. The samples were stored in in 100% MeOH at −20°C overnight.

The samples were rehydrated using sequential 5-minute washes in 75%, 50%, and 25% MeOH:PBS solutions while rocking and on ice, followed by two 5-minute washes in PBS, rocking and on ice. Proteinase K (PK) solution was applied to the samples at a dilution of 1:2000 in PK:PBS for time t (t = 15 seconds * HH stage). The samples were postfixed in 4% PFA for 20 minutes at room temperature and underwent 2 x 5 minute PBS washes, followed by 50% 5x Standard Saline Citrate with Tween (SSCT) and a final wash in 5x SSCT. HCR v3.0 was performed using the protocol as described by Molecular Instruments (Choi et al., 2018). Split initiator probes (v3.0) for HMGA1 (accession NM_204369.2), KPNA2 (accession NM_001006209.2) and CSRP2 (accession NM_205208.2) were designed by Molecular Instruments.

For imaging on the Zeiss Lightsheet 7 samples were first cleared in SCALE A2 at 4°C overnight.

### Statistical analysis

Limbs were imaged on the Leica MZ8 light microscope and anatomical lengths were measured using ImageJ (Schindelin et al., 2012). Comparison between the manipulated and contralateral digits (in both the experimental and control groups) were carried out using a paired T test. Comparisons between experimental and control groups were performed using an unpaired T test.

### mRNA seq analysis

#### Raw read alignment and quantification

Single cell suspensions were sequenced by the Tschopp lab using the 10X Genomics Chromium Single Cell System, with a suspension concentration of approximately 4000 cells microliter^-1^. The raw FASTQ reads were pre-processed using Trim Galore to remove adapter sequences from the raw sequencing reads, and to trim low quality reads that occur at the end of the reads. The processed reads were aligned to the ENSEMBL chicken genome (GRCg7b) assembly and quantified by STARsolo.

### Quality control

The generated cell and count matrices were imported into R and converted into Seurat objects using the Seurat package (Stuart et al., 2019), encompassing 7,168 and 5,672 cells for the HH 25 and HH 31 datasets, respectively. For compatibility with other analytical tools, these Seurat objects were subsequently converted into SingleCellExperiment objects (Amezquita et al., 2020).

Each dataset was subjected to an independent quality control (QC) pipeline prior to integration. Outlier cells were identified using the *isOutlier* function in the *scater* package (McCarthy et al., 2017), applying median absolute deviation thresholds of 3, 3, and 5 for the number of counts, number of features, and mitochondrial gene expression percentage, respectively. Ambient RNA contamination was detected and removed using *DecontX* from the *celda* package (Yang et al., 2020), and doublets were filtered out using *ScDblFinder* (Germain et al., 2021).

Cell cycle scoring was performed using an updated 2019 gene set (Stuart et al., 2019) and regressed out using *SCTransform*. After initial QC, the two datasets were integrated, and a second QC was applied to the integrated object, removing any remaining outliers or low-quality cells.

The integrated object contained 6,769 and 5,438 cells from the HH 25 and HH 31 datasets respectively.

### Cluster generation and annotation

Clustering was performed on the integrated dataset through RunPCA, FindNeighbors and FindClusters. FindNeighbors was run using the first PC that had a cumulative percent greater than 90% and <5% associated variation. To find marker genes for each cluster differential expression analysis was performed using FindMarkers (test.use = “MAST”).

### Trajectory analysis

The trajectory analysis was carried out using *Slingshot* (Street et al., 2018), where cluster “5” and “1” were set as the starting and ending cluster respectively.

## Supporting information

Supplemental Information 2 - Differential expression

Supplemental Information 3 - Comparative statistics

## Figures

All figures were made using BioRender.com

## Acknowledgements

We thank the National Avian Research Facility, The Roslin Institute and R(D)SVS for the maintenance and production of fertile eggs.

## Author Contributions

CBS undertook experimental design, experiments, analysis, manuscript preparation, JS undertook experimental design, experiments, analysis. LK undertook experimental design, experiments, analysis. NC undertook experimental design, experiments, analysis. LM undertook technical support. MD conceived the study, undertook experimental design, experiments, analysis, manuscript preparation.

During the preparation of this work the author(s) used ChatGPT in order to help proof read. After using this tool/service, the author(s) reviewed and edited the content as needed and take(s) full responsibility for the content of the publication.

## References

Algaze, I., Snyder, A. J., Hodges, N. L., & Smith, G. A. (2012). Children Treated in United States Emergency Departments for Door-Related Injuries, 1999-2008. Clinical pediatrics, 51(3), 226–232. 10.1177/0009922811423308

Amezquita, R. A., Lun, A. T. L., Becht, E., Carey, V. J., Carpp, L. N., Geistlinger, L.,…Hicks, S. C. (2020). Orchestrating single-cell analysis with Bioconductor. Nature methods, 17(2), 137–145. 10.1038/s41592-019-0654-x

Bermudez VP, Maniwa Y, Tappin I, Ozato K, Yokomori K, Hurwitz J. The alternative Ctf18-Dcc1-Ctf8-replication factor C complex required for sister chromatid cohesion loads proliferating cell nuclear antigen onto DNA. (2003) PNAS.;100(18):10237–42. doi: 10.1073/pnas.1434308100. Epub 2003 Aug 20. PMID: 12930902; PMCID: PMC193545.

Bi, W., Deng, J. M., Zhang, Z., Behringer, R. R., & De Crombrugghe, B. (1999). Sox9 is required for cartilage formation. Nature Genetics, 22(1), 85–89. 10.1038/8792

Caplan, A. I., & Koutroupas, S. (1973). The control of muscle and cartilage development in the chick limb: the role of differential vascularization. Journal of embryology and experimental morphology, 29(3), 571–583. 10.1242/dev.29.3.571

Choi, H. M. T., Schwarzkopf, M., Fornace, M. E., Acharya, A., Artavanis, G., Stegmaier, J.,…Pierce, N. A. (2018). Third-generation in situ hybridization chain reaction: multiplexed, quantitative, sensitive, versatile, robust. Development, 145(12), dev165753–dev165753. 10.1242/dev.165753

Cooper, L. N., Berta, A., Dawson, S. D., & Reidenberg, J. S. (2007). Evolution of hyperphalangy and digit reduction in the cetacean manus. Anatomical record (Hoboken, N.J. : 2007), 290(6), 654–672. 10.1002/ar.20532

Dahn, R. D., & Fallon, J. F. (2000). Interdigital Regulation of Digit Identity and Homeotic Transformation by Modulated BMP Signaling. Science, 289(5478), 438–441. 10.1126/science.289.5478.438

Davey, M. G., Towers, M., Vargesson, N., & Tickle, C. (2018). The chick limb: embryology, genetics and teratology. The International journal of developmental biology, 62(1-2-3), 85-95. 10.1387/ijdb.170315CT

Feinberg, R. N., & Beebe, D. C. (1983). Hyaluronate in Vasculogenesis. Science., 220(4602), 1177–1179. 10.1126/science.6857242

Feregrino C, Sacher F, Parnas O, Tschopp P (2019). A single-cell transcriptomic atlas of the developing chicken limb. BMC Genomics;20(1):401. doi: 10.1186/s12864-019-5802-2. PMID: 31117954; PMCID: PMC6530069.

Gandhi, S., Hutchins, E. J., Maruszko, K., Park, J. H., Thomson, M., & Bronner, M. E. (2020). Bimodal function of chromatin remodeler hmga1 in neural crest induction and wnt-dependent emigration. eLife, 9, 1–62. 10.7554/ELIFE.57779

Germain, P.-L., Lun, A., Garcia Meixide, C., Macnair, W., & Robinson, M. D. (2021). Doublet identification in single-cell sequencing data using scDblFinder F1000 research, 10, 979–979. 10.12688/f1000research.73600.2

Grall E, Feregrino C, Fischer S, De Courten A, Sacher F, Hiscock TW, Tschopp P (2024). Self-organized BMP signaling dynamics underlie the development and evolution of digit segmentation patterns in birds and mammals. PNAS;121(2):e2304470121. doi: 10.1073/pnas.2304470121. Epub 2024 Jan 4. PMID: 38175868; PMCID: PMC10786279.

Hamburger V, Hamilton HL. A series of normal stages in the development of the chick embryo. J Morphol. 1951 Jan;88(1):49–92. PMID: 24539719.

Hiscock, T. W., Tschopp, P., & Tabin, C. J. (2017). On the Formation of Digits and Joints during Limb Development. Developmental Cell, 41(5), 459–465. 10.1016/j.devcel.2017.04.021

Huang, B.-L., Trofka, A., Furusawa, A., Norrie, J. L., Rabinowitz, A. H., Vokes, S. A.,…Mackem, S. (2016). An interdigit signalling centre instructs coordinate phalanx-joint formation governed by 5′Hoxd–Gli3 antagonism. Nature Communications, 7(1), 12903. 10.1038/ncomms12903

Kaplan, D. R., & Miller, F. D. (2000). Neurotrophin signal transduction in the nervous system. Current Opinion in Neurobiology, 10(3), 381–391. 10.1016/S0959-4388(00)00092-1

Lefebvre, V., & Smits, P. (2005). Transcriptional control of chondrocyte fate and differentiation. Birth Defects Research Part C: Embryo Today: Reviews, 75(3), 200–212. 10.1002/bdrc.20048

Lin, X., Mekonnen, T., Verma, S., Zevallos-Delgado, C., Singh, M., Aglyamov, S. R.,…Coulson-Thomas, V. J. (2022). Hyaluronan Modulates the Biomechanical Properties of the Cornea. Investigative Opthalmology & Visual Science, 63(13), 6. 10.1167/iovs.63.13.6

Liu, C., Lin, C., Gao, C., May-Simera, H., Swaroop, A., & Li, T. (2014). Null and hypomorph *Prickle1*alleles in mice phenocopy human Robinow syndrome and disrupt signaling downstream of Wnt5a. Biology Open, 3(9), 861–870. 10.1242/bio.20148375

McCarthy, D. J., Campbell, K. R., Lun, A. T. L., & Wills, Q. F. (2017). Scater: pre-processing, quality control, normalization and visualization of single-cell RNA-seq data in R. Bioinformatics, 33(8), 1179–1186. 10.1093/bioinformatics/btw777

Mcclean, D., & Rogers, H. J. (1945). Structure of Wharton’s Jelly. Nature, 155(3942), 606–607. 10.1038/155606b0

Montero, J. A., Lorda-Diez, C. I., Gañan, Y., Macias, D., & Hurle, J. M. (2008). Activin/TGFβ and BMP crosstalk determines digit chondrogenesis. Developmental biology, 321(2), 343–356. 10.1016/j.ydbio.2008.06.022

Norrie JL, Li Q, Co S, Huang BL, Ding D, Uy JC, Ji Z, Mackem S, Bedford MT, Galli A, Ji H, Vokes SA(2016). PRMT5 is essential for the maintenance of chondrogenic progenitor cells in the limb bud. Development;143(24):4608–4619. doi: 10.1242/dev.140715. Epub 2016 Nov 8. PMID: 27827819; PMCID: PMC5201029.

Nurse P. Universal control mechanism regulating onset of M-phase. (1990). Nature;344(6266):503–8. doi: 10.1038/344503a0. PMID: 2138713.

Mui B.W.H., Wong J.Y., Bray T, Connolly L, Wang J.H., Winkel A., Robey P.G., Franze K., Chalut K.J., Storer M.A.(2024) Hyaluronic Acid and Emergent Tissue Mechanics Orchestrate Digit Tip Regeneration. bioRxiv 2024.12.04.626830; doi: 10.1101/2024.12.04.626830

Oh, J. D. H., Freem, L., Saunders, D. D. Z., Mcteir, L., Gilhooley, H., Jackson, M.,…Davey, M. G. (2024). Insights into digit evolution from a fate map study of the forearm using Chameleon, a new transgenic chicken line. Development, 151(13). 10.1242/dev.202340

Parada C, Banavar SP, Khalilian P, Rigaud S, Michaut A, Liu Y, Joshy DM, Campàs O, Gros J. (2022). Mechanical feedback defines organizing centers to drive digit emergence. Dev Cell. 11;57(7):854–866.e6. doi: 10.1016/j.devcel.2022.03.004. PMID: 35413235.

Reid, D. B. C., Shah, K. N., Eltorai, A. E. M., Got, C. C., & Daniels, A. H. (2019). Epidemiology of Finger Amputations in the United States From 1997 to 2016. Journal of hand surgery global online, 1(2), 45–51. 10.1016/j.jhsg.2019.02.001

Richardson, M. K., & Oelschläger, H. H. A. (2002). Time, pattern, and heterochrony: a study of hyperphalangy in the dolphin embryo flipper. Evolution & development, 4(6), 435–444. 10.1046/j.1525-142X.2002.02032.x

Sanz-Ezquerro JJ, Tickle C. (2003) Fgf signaling controls the number of phalanges and tip formation in developing digits. Curr Biol. (20):1830–6. doi: 10.1016/j.cub.2003.09.040. PMID: 14561411.

Saxena, A., Towers, M., & Cooper, K. L. (2017). The origins, scaling and loss of tetrapod digits. PNAS, 372(1713), 20150482. 10.1098/rstb.2015.0482

Schindelin J, Arganda-Carreras I, Frise E, Kaynig V, Longair M, Pietzsch T, Preibisch S, Rueden C, Saalfeld S, Schmid B, Tinevez JY, White DJ, Hartenstein V, Eliceiri K, Tomancak P, Cardona A.(2012). Fiji: an open-source platform for biological-image analysis. Nat Methods;9(7):676–82. doi: 10.1038/nmeth.2019. PMID: 22743772; PMCID: PMC3855844.

Scoones, J. C., & Hiscock, T. W. (2020). A dot-stripe Turing model of joint patterning in the tetrapod limb. Development, 147(8). 10.1242/dev.183699

Seger, R., & Krebs, E. G. (1995). The MAPK signaling cascade. The FASEB journal, 9(9), 726–735. 10.1096/fasebj.9.9.7601337

Sensiate, L. A., & Marques-Souza, H. (2019). Bone growth as the main determinant of mouse digit tip regeneration after amputation. Scientific reports, 9(1), 9720–9728. 10.1038/s41598-019-45521-4

Sobolewski, K., Małkowski, A., Bańkowski, E., & Jaworski, S. (2005). Wharton’s jelly as a reservoir of peptide growth factors. Placenta (Eastbourne), 26(10), 747–752. 10.1016/j.placenta.2004.10.008

Storm, E. E., & Kingsley, D. M. (1999). GDF5 Coordinates Bone and Joint Formation during Digit Development. Developmental biology., 209(1), 11–27. 10.1006/dbio.1999.9241

Street, K., Risso, D., Fletcher, R. B., Das, D., Ngai, J., Yosef, N.,…Dudoit, S. (2018). Slingshot: cell lineage and pseudotime inference for single-cell transcriptomics. BMC genomics, 19(1), 477–477. 10.1186/s12864-018-4772-0

Stricker, S., & Mundlos, S. (2011). Mechanisms of digit formation: Human malformation syndromes tell the story. Developmental dynamics, 240(5), 990–1004. 10.1002/dvdy.22565

Stuart, T., Butler, A., Hoffman, P., Hafemeister, C., Papalexi, E., Mauck, W. M.,…Satija, R. (2019). Comprehensive Integration of Single-Cell Data. Cell, 177(7), 1888–1902.e1821. 10.1016/j.cell.2019.05.031

Sun, X., Martin, G. R., & Lewandoski, M. (2000). Fgf8 signalling from the AER is essential for normal limb development. Nature genetics, 26(4), 460–463. 10.1038/82609

Suzuki, T., Hasso, S. M., & Fallon, J. F. (2008). Unique SMAD1/5/8 activity at the phalanx-forming region determines digit identity. Proceedings of the National Academy of Sciences - PNAS, 105(11), 4185–4190. 10.1073/pnas.0707899105

Temtamy, S. A., & Aglan, M. S. (2008). Brachydactyly. Orphanet Journal of Rare Diseases, 3(1), 15. 10.1186/1750-1172-3-15

Tiecke, E., & Tickle, C. (2007). Application of sonic hedgehog to the developing chick limb. *Methods in molecular biology (Clifton*, N.J.), 397, 23. 10.1385/1-59745-516-4:23

Ullrich, A., & Schlessinger, J. (1990). Signal transduction by receptors with tyrosine kinase activity. Cell, 61(2), 203–212. 10.1016/0092-8674(90)90801-K

Witte, F., Chan, D., Economides, A. N., Mundlos, S., & Stricker, S. (2010). Receptor tyrosine kinase-like orphan receptor 2 (ROR2) and Indian hedgehog regulate digit outgrowth mediated by the phalanx-forming region. Proceedings of the National Academy of Sciences, 107(32), 14211–14216. 10.1073/pnas.1009314107

Yang, S., Corbett, S. E., Koga, Y., Wang, Z., Johnson, W. E., Yajima, M., & Campbell, J. D. (2020). Decontamination of ambient RNA in single-cell RNA-seq with DecontX. Genome Biology, 21(1), 1–57. 10.1186/s13059-020-1950-6

Yang, T., Bassuk, A. G., & Fritzsch, B. (2013). Prickle1 stunts limb growth through alteration of cell polarity and gene expression. Developmental Dynamics, 242(11), 1293–1306. 10.1002/dvdy.24025

Zhu Z, Liu Y, Zeng J, Ren S, Wei L, Wang F, Sun X, Huang Y, Jiang H, Sui X, Jin W, Jin L, Sun X. Diosbulbin C, a novel active ingredient in Dioscorea bulbifera L. extract, inhibits lung cancer cell proliferation by inducing G0/G1 phase cell cycle arrest. (2023). BMC Complement Med Ther.23(1):436. doi: 10.1186/s12906-023-04245-9. PMID: 38049779; PMCID: PMC10694954.

