## Supplemental Information 1 - Additional figures for "Characterisation of the Avascular Mesenchyme during Digit Outgrowth"

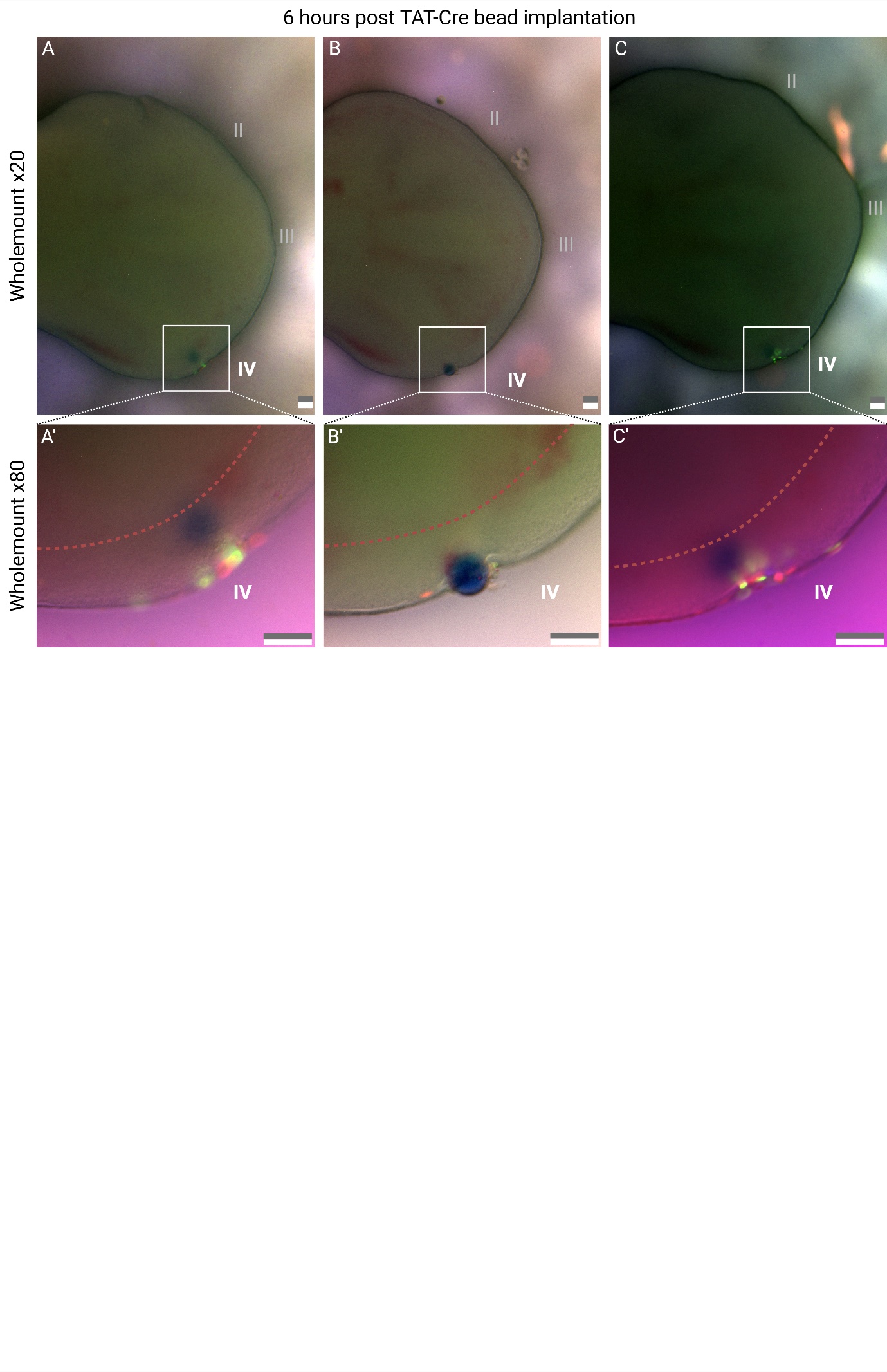

**Figure S1.** **Fluorescent labelling of cells in proximity to** **Affi-Gel Blue Gel beads soaked in TAT-Cre**. Embryos were manipulated *in ovo*, whereby an Affi-Gel Blue Gel bead containing TAT-Cre was inserted into digit IV and embryos were resealed and incubated at 38°C. (**A, B, C**) Wholemount fluorescent images of CPX embryo hindlimbs 6 hours post bead insertion. Regions outlined by white boxes are enlarged below (**A’, B’, C’**), which demonstrated that only cells in close proximity to the beads are labelled. Of note, no labelled cells were identified proximal to the avascular mesenchyme (proximal boundary demarked by red dotted line), with labelled cells only observed within the distal ectoderm and avascular mesenchyme. As labelling of cells occurs immediately and very transiently the fluorescent cells in **Figure 2 I-J** represent the clones of cells from the ectoderm and avascular mesenchyme exclusively. Scale bars = 100 µm.

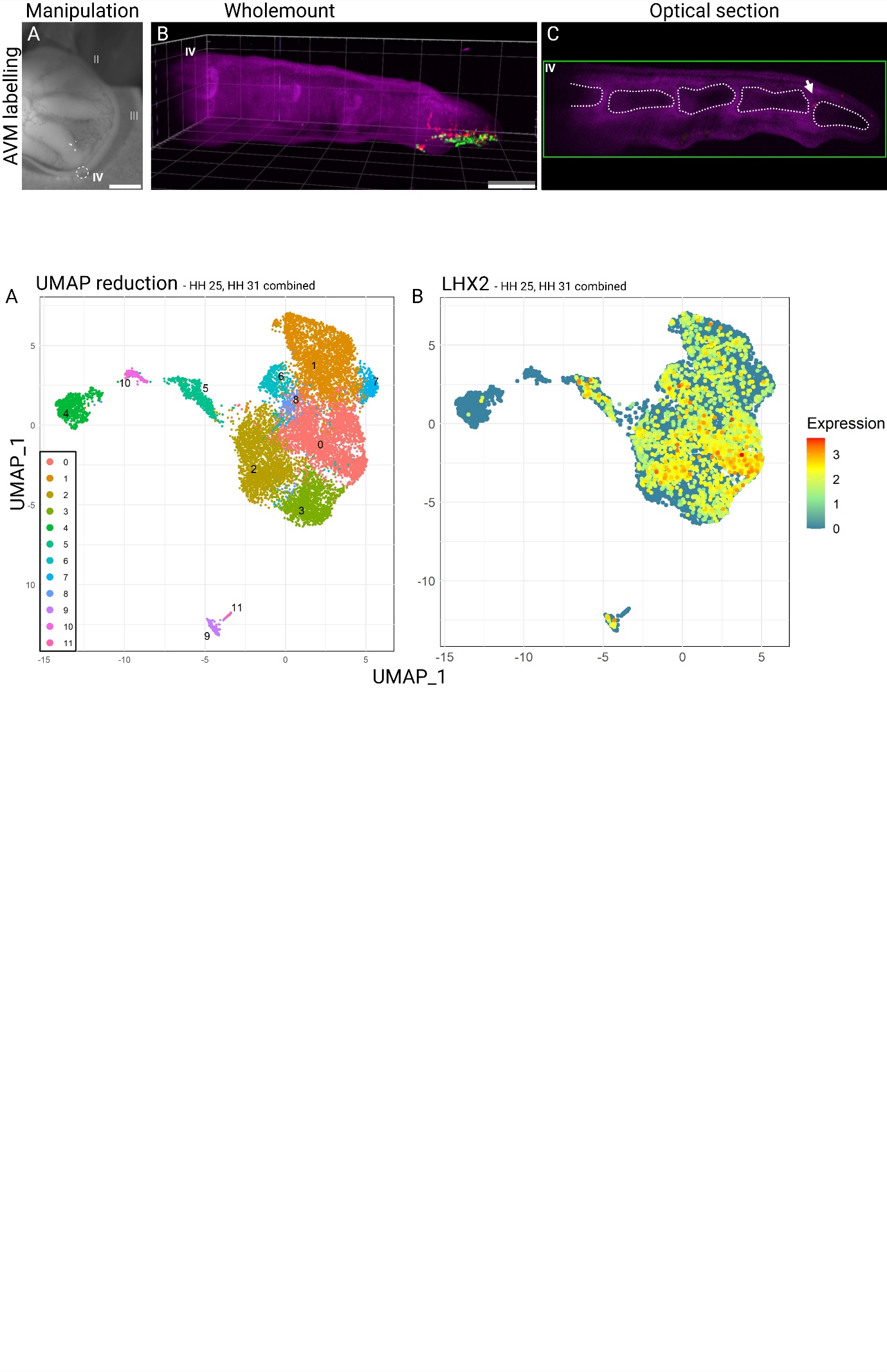

**Figure S2.** **Sparse labelling of the avascular mesenchyme distal to digit IV**. (**A**) Beads soaked in TAT-Cre were inserted at the indicated sites (white dotted circles) in HH 27 hindlimb autopods and left for 60 seconds before being removed. (**B**) Lightsheet wholemount fluorescent image showing clones of labelled cells from the Avascular Mesenchyme (AVM) in **A**. (**C**) Optical section through digit IV, with a white arrow indicating labelled cells located within the synovial joint between phalanx 4 and the terminal phalanx. These cells accounted for a small percentage of the total number of labelled cells. Scale bar = 500 µm. **Related to The Avascular Mesenchyme is not determined, the Phalanx Forming Region is determined.**

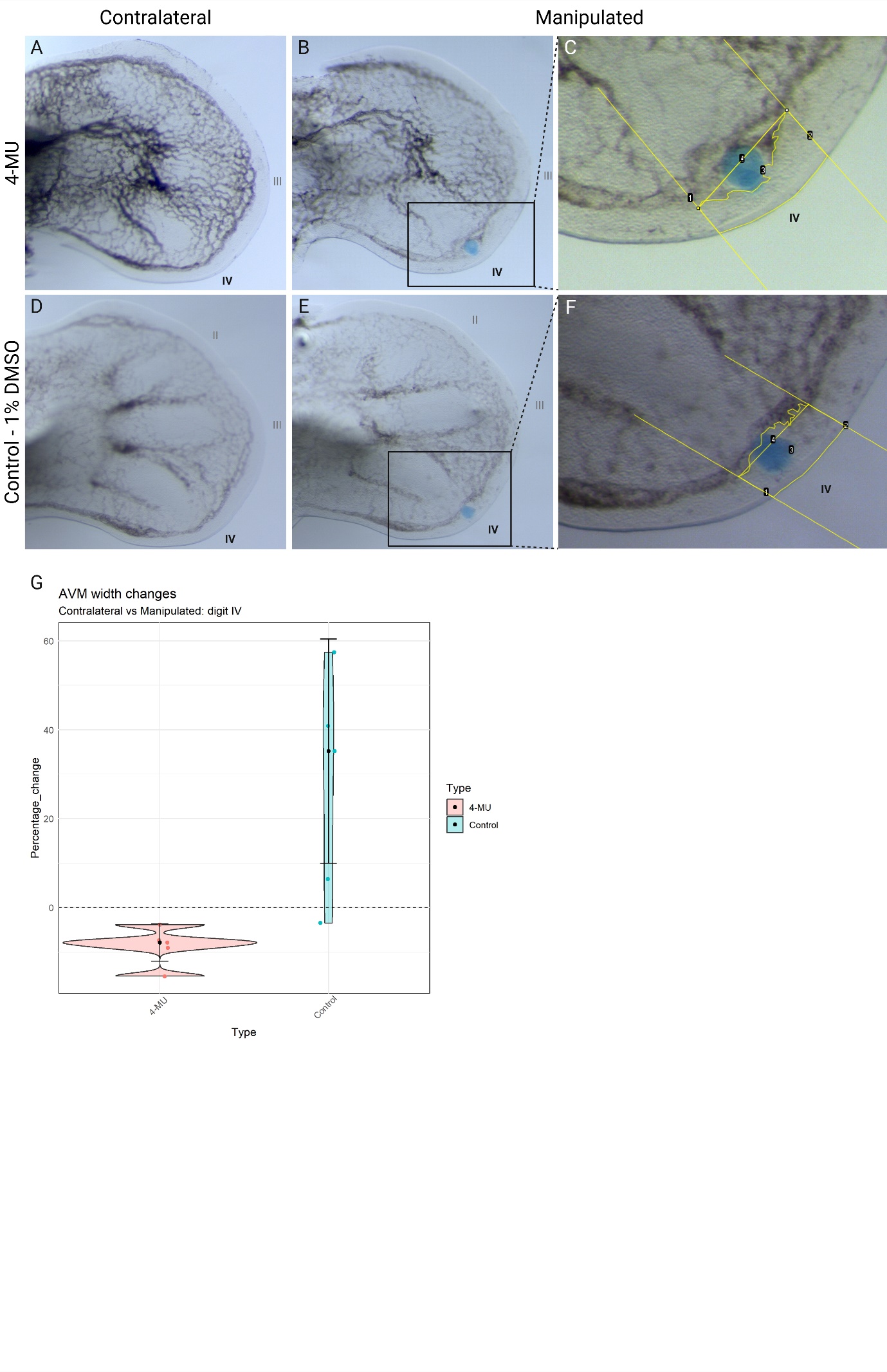

**Figure S3.** **Inhibition of hyaluronic acid synthesis leads to local reductions in the width of the avascular mesenchyme**. (**A-F**) Vascular India ink injection of an HH 28 chick hindlimb autopod 6 hours post insertion into digit IV of a Affi-Gel Blue Gel bead soaked in 0.1 mM of hyaluronic acid synthesis inhibitor 4-methylumbelliferone (4-MU) (1% DMSO) (**B**) or a control solution of DMEM (1% DMSO) (**E**) (n = 5, n =5 respectively). Regions outlined by black boxes lines are enlarged to the right (**C, F**) where the total area of the avascular mesenchyme distal to digit IV was measured in both the experimental and control (n = 5, n = 5 for experimental and control respectively). Yellow lines indicate the measured area of the avascular mesenchyme (line 3). Percentage difference in normalised area was assessed through comparison of each manipulated limb to its corresponding contralateral region. (**G**) Violin plot comparing the percentage change in normalised avascular mesenchyme area distal to digit IV. Analysis identified a significant reduction (p = 0.014) in the normalised area of the avascular mesenchyme when treated with 4-MU. **Related to Ectodermal signalling to the Avascular Mesenchyme is essential for proper phalangeal patterning.**

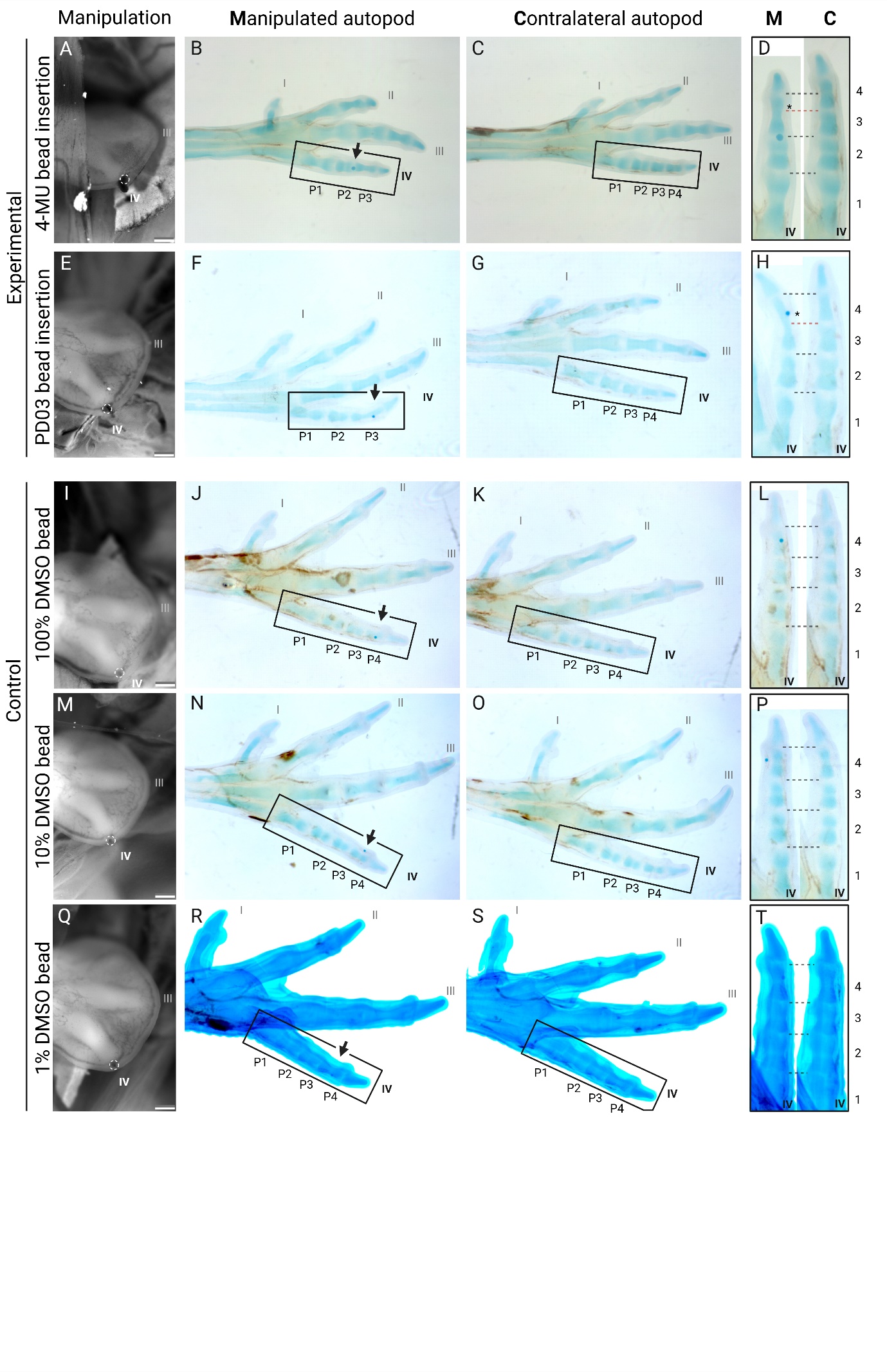

**Figure S4.** **Control bead implantation into the avascular mesenchyme does not cause phalangeal abnormalities.** (**A, E**) Beads soaked in 4-MU or PD0325901 were implanted distal to digit IV in HH27 embryos. White dotted circles indicate bead locations. (**B-D, F-H**) At embryonic day (E) 11 cartilage staining and clearing revealed that the manipulated digits lacked phalanx (P4) and exhibited significant elongation of P3 compared to the contralateral digits. Black arrows mark the original bead site, and asterisks indicate expected synovial joint locations. (**I-T**) Control beads soaked in DMSO (vehicle) were implanted in identical regions as the experimental group in HH 27 embryos (n = 3 for each DMSO concentration respectively). No skeletal abnormalities were observed at any DMSO concentration, confirming that the defects in **Figure 4** resulted from the pharmacological agents. Scale bars = 500 µm.

| Cluster(s) | Cell type | Marker genes | Reference |
| --- | --- | --- | --- |
| 2, 3, 7 | Mesenchyme | PRRX1 | (Kuratani et al., 1994) |
| 0 | Distal mesenchyme | PCP4, MSX1 | (Feregrino et al., 2019) (Yokouchi et al., 1991) |
| 6, 8 | Interdigital mesenchyme | RDH10, BMP4 | (Cunningham et al., 2011) (Cammas et al., 2007) (Macias et al., 1997) |
| 5 | Avascular mesenchyme | PRMT5 | (Norrie et al., 2016) |
| 1 | Chondrocytes | SOX9, ACAN | (Lefebvre & Smits, 2005) (Bi et al., 1999) |
| 4 | Epithelium | KRT14, CDH3 | (Wang et al., 2020) (Blanpain & Fuchs, 2009) |
| 9, 11 | Erythrocyte | HBA1, HBAD | (Perutz & Lehmann, 1968) |
| 10 | Unknown |  |  |

**Table S1.** Predicted cell types, enriched marker genes and references for the annotation of clusters in **Figure 5**.

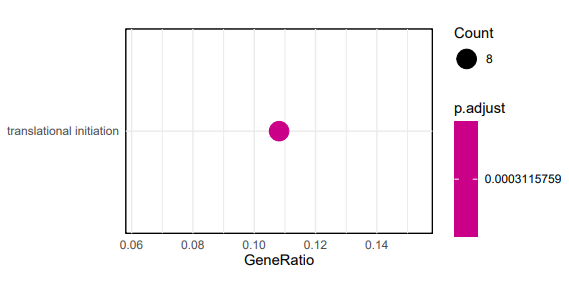

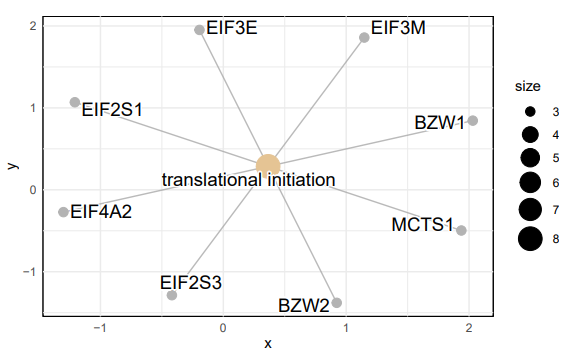

**Figure S5.** Gene set enrichment output of top 200 enriched and differentially expressed genes identified in cluster 5. **Related to Transcriptional profiling of the Avascular Mesenchyme.**

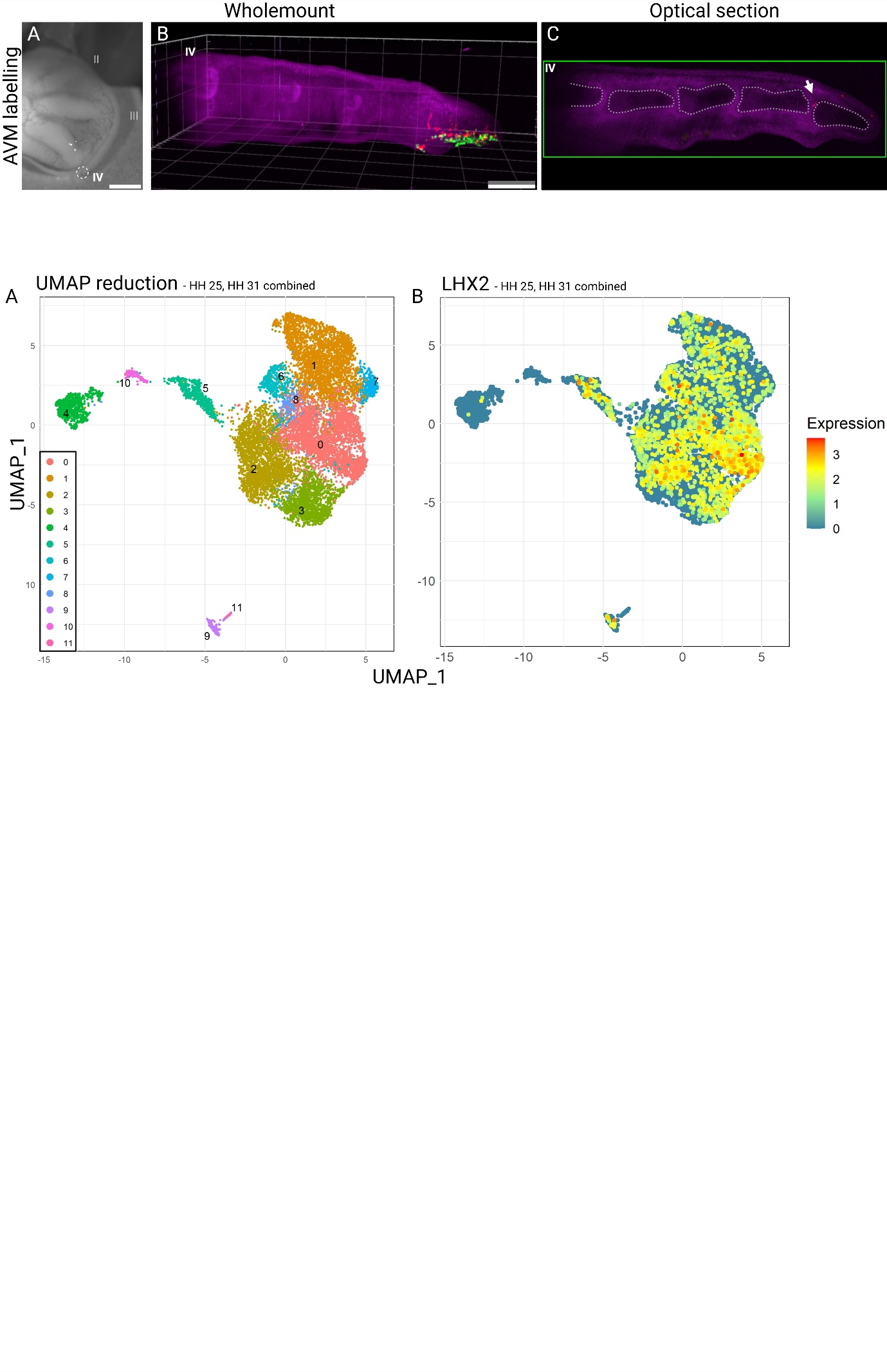

**Figure S6.** **Expression of *LHX2* identified within the single cell mRNA-sequencing data**. (**A**) UMAP representation of the combined HH 25 and HH 31 datasets (Feregrino et al., 2019). (**B**) Feature plot showing the expression of the distal mesenchyme marker gene *LHX2*. It features expression within the distal mesenchyme cluster (cluster 0) and the avascular mesenchyme enriched cluster (cluster 5). *LHX2* expression was not differentially expressed within cluster 5. **Related to Transcriptional profiling of the Avascular Mesenchyme.**

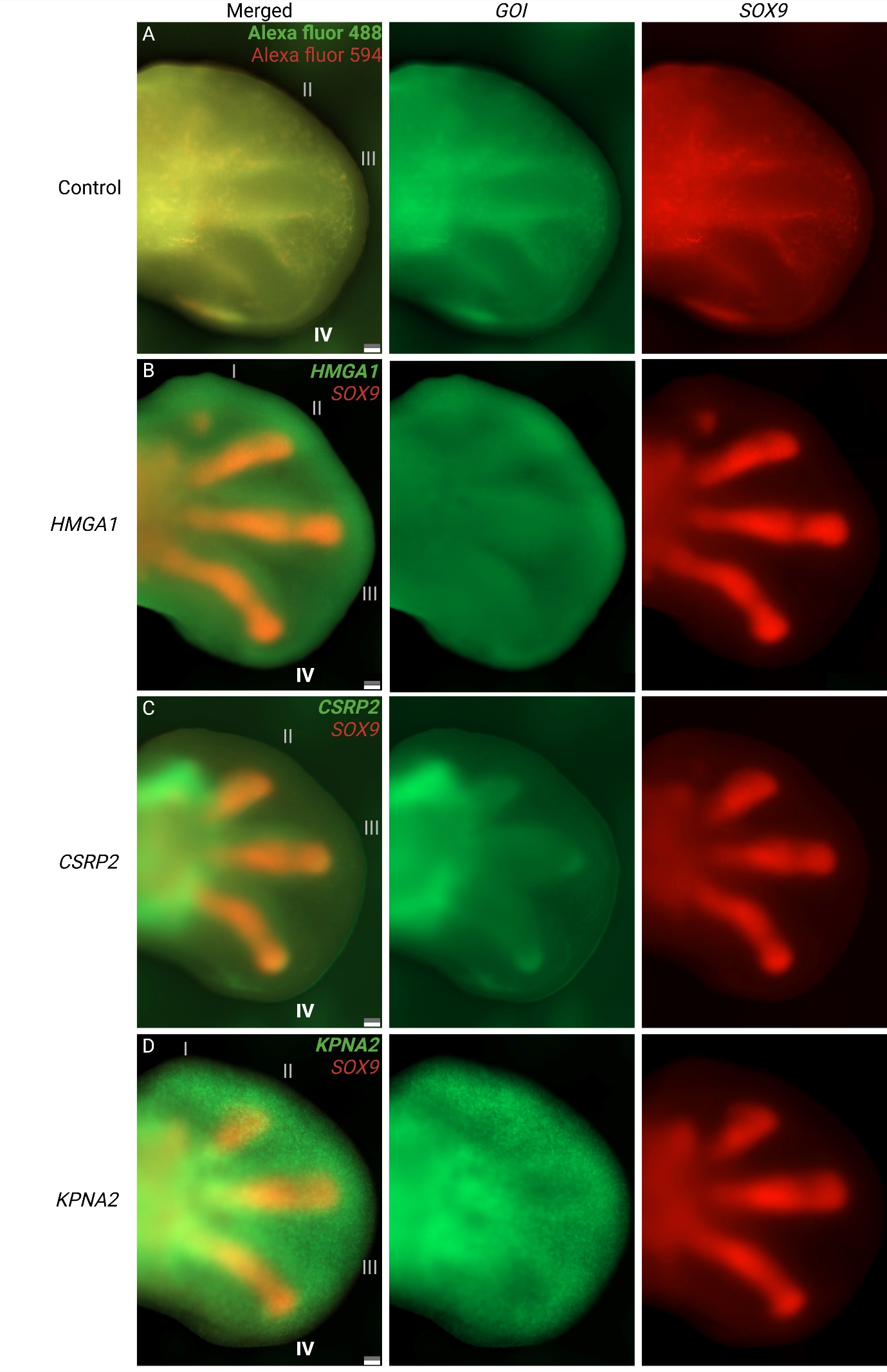

**Figure S7.** **Fluorescent wholemount images of differentially expressed genes in cluster 5.** *In situ* hybridisation images of genes identified as significantly positively differentially expressed within cluster 5 (enriched with cells from the avascular mesenchyme). (**A**) Control wholemount images using the same fluorescent hairpins as in (**B-D**) but without a gene-specific probe, allowing visualisation of non-specific hairpin binding. While broad non-specific binding and autofluorescence were observed, the control exhibited fluorescence patterns distinct from the specific binding seen in (**B-D**). Notably, the avascular mesenchyme, lacking blood vessels, displayed lower fluorescence intensity than the more proximal regions, supporting that the fluorescent wholemount images in **Figure 5** represent genuine gene expression patterns. Scale bars = 100 µm.

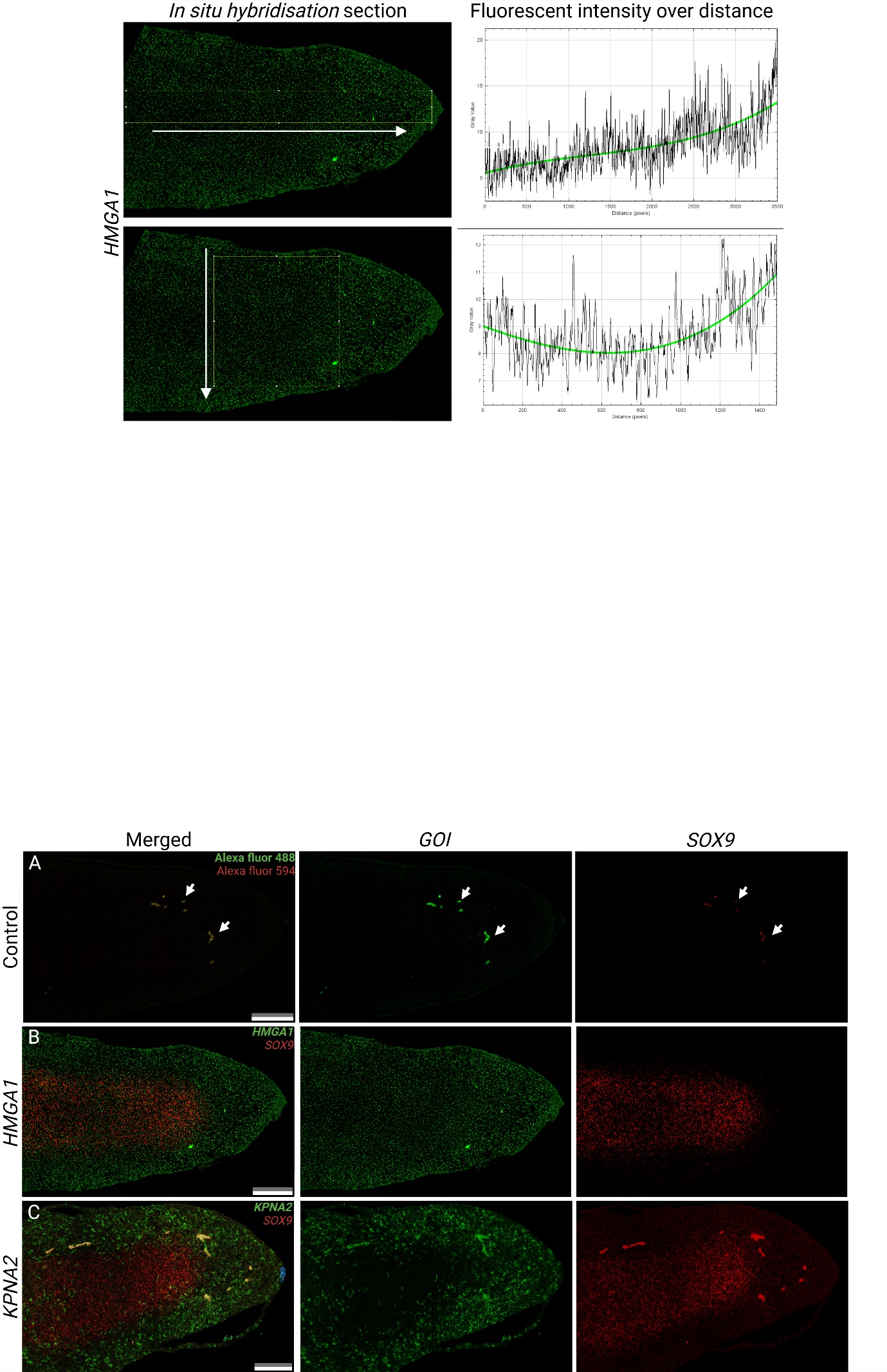

Figure S8. **Fluorescent images of sections of differentially expressed genes in cluster 5.** *In situ* hybridisation images of genes identified as significantly positively differentially expressed within cluster 5 (enriched with cells from the avascular mesenchyme). (**A**) Control images of sections using the same fluorescent hairpins as in (**B-C**) but without a gene-specific probe, allowing visualisation of non-specific hairpin binding. Very low levels of non-specific binding were observed, and autofluorescent blood vessels are denoted by white arrows. This supports that the fluorescent images of sections in **Figure 5** represent genuine gene expression patterns. Scale bars = 100 µm.

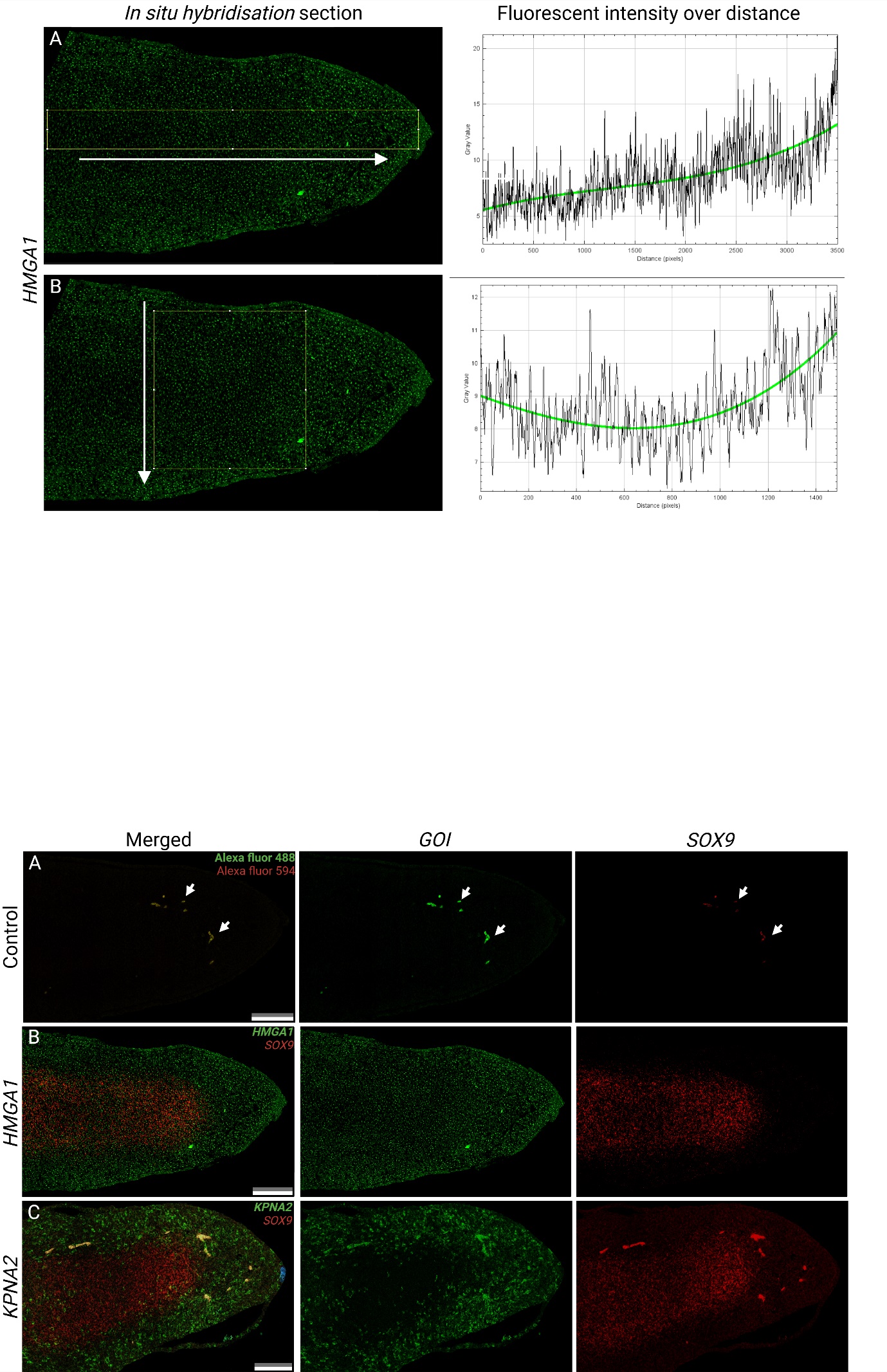

**Figure S9.** **Patterns of *HMGA1* expression within a sagittal section**. Analysis of fluorescent *in situ* hybridisation sections of *HMAG1* revealed changes in fluorescent intensity across the regions of interest (area constrained by yellow boxes). (**A**) Increasing levels of *HMGA1* expression along the proximal-distal axis (white arrow), with greatest expression within the distal aspect of the digit – region contained by the avascular mesenchyme. (**B**) Elevated expression of *HMGA1* within the more superficial digit regions across the dorsal-ventral axis (white arrow). **Related to Transcriptional profiling of the Avascular Mesenchyme.**

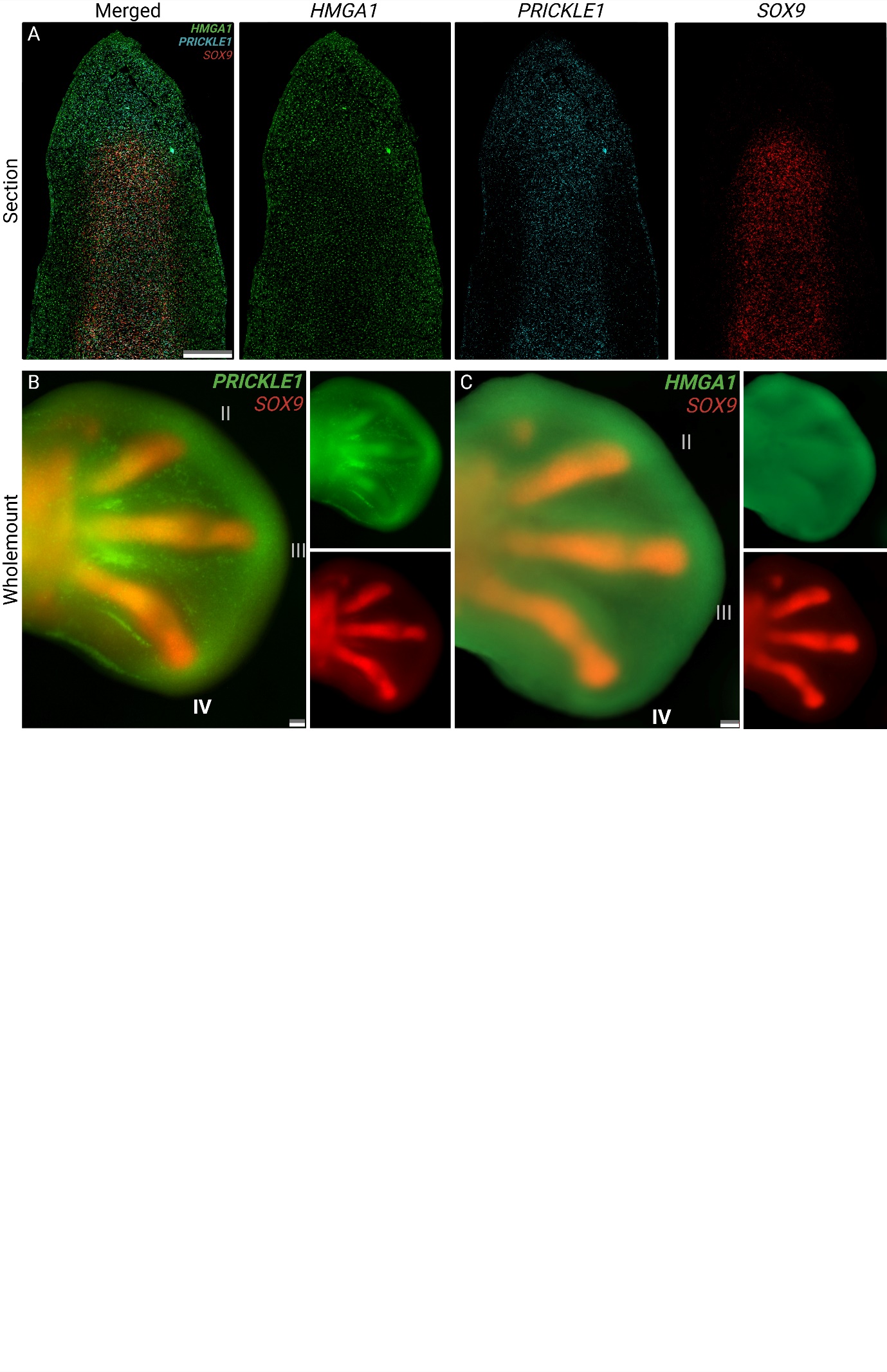

**Figure S10.** ***HMGA1* and *PRICKLE* display highly similar expression domains**. (**A**) Multiplexed *in situ* hybridisation sagittal section of a digit from a HH27 hindlimb. Both *HMGA1* and *PRICKLE1* show similar patterns of elevated expression within the distal and avascular mesenchyme. *PRICKLE*, however, also exhibits prominent expression within the digital ray as indicted by its overlapping expression domain with *SOX9*. (**B**) *In situ hybridisation* wholemount of *PRICKLE1* expression in a HH27 hindlimb. Of note, *PRICKLE1* displays elevated expression within the distal and avascular mesenchyme and is focally elevated distal to the digits. (**C**) *In situ* hybridisation wholemount of *HMAG1* expression in a HH27 hindlimb. *HMGA1* shares the same expression characteristics as *PRICKLE1* within the distal and avascular mesenchyme. Scale bars = 100 µm. **Related to Transcriptional profiling of the Avascular Mesenchyme.**

**Supplemental references:**

Bi, W., Deng, J. M., Zhang, Z., Behringer, R. R., & De Crombrugghe, B. (1999). Sox9 is required for cartilage formation. *Nature Genetics*, *22*(1), 85-89. <https://doi.org/10.1038/8792>

Blanpain, C., & Fuchs, E. (2009). Epidermal homeostasis: a balancing act of stem cells in the skin. *Nature reviews. Molecular cell biology*, *10*(3), 207-217. <https://doi.org/10.1038/nrm2636>

Cammas, L., Romand, R., Fraulob, V., Mura, C., & Dollé, P. (2007). Expression of the murine retinol dehydrogenase 10 (Rdh10) gene correlates with many sites of retinoid signalling during embryogenesis and organ differentiation. *Developmental dynamics*, *236*(10), 2899-2908. <https://doi.org/10.1002/dvdy.21312>

Cunningham, T. J., Chatzi, C., Sandell, L. L., Trainor, P. A., & Duester, G. (2011). Rdh10 mutants deficient in limb field retinoic acid signaling exhibit normal limb patterning but display interdigital webbing. *Developmental dynamics*, *240*(5), 1142-1150. <https://doi.org/10.1002/dvdy.22583>

Feregrino, C., Sacher, F., Parnas, O., & Tschopp, P. (2019). A single-cell transcriptomic atlas of the developing chicken limb. *BMC Genomics*, *20*(1). <https://doi.org/10.1186/s12864-019-5802-2>

Kuratani, S., Martin, J. F., Wawersik, S., Lilly, B., Eichele, G., & Olson, E. N. (1994). The Expression Pattern of the Chick Homeobox Gene gMHox Suggests a Role in Patterning of the Limbs and Face and in Compartmentalization of Somites. *Developmental biology*, *161*(2), 357-369. <https://doi.org/10.1006/dbio.1994.1037>

Lefebvre, V., & Smits, P. (2005). Transcriptional control of chondrocyte fate and differentiation. *Birth Defects Research Part C: Embryo Today: Reviews*, *75*(3), 200-212. <https://doi.org/10.1002/bdrc.20048>

Macias, D., Gañan, Y., Sampath, T. K., Piedra, M. E., Ros, M. A., & Hurle, J. M. (1997). Role of BMP-2 and OP-1 (BMP-7) in programmed cell death and skeletogenesis during chick limb development. *Development*, *124*(6), 1109-1117. <https://doi.org/10.1242/dev.124.6.1109>

Norrie, J. L., Li, Q., Co, S., Huang, B.-L., Ding, D., Uy, J. C.,…Vokes, S. A. (2016). PRMT5 is essential for the maintenance of chondrogenic progenitor cells in the limb bud. *Development*, *143*(24), 4608-4619. <https://doi.org/10.1242/dev.140715>

Perutz, M. F., & Lehmann, H. (1968). Molecular Pathology of Human Haemoglobin. *Nature (London)*, *219*(5157), 902-909. <https://doi.org/10.1038/219902a0>

Wang, S., Drummond, M. L., Guerrero-Juarez, C. F., Tarapore, E., MacLean, A. L., Stabell, A. R.,…Atwood, S. X. (2020). Single cell transcriptomics of human epidermis identifies basal stem cell transition states. *Nature communications*, *11*(1), 4239-4239. <https://doi.org/10.1038/s41467-020-18075-7>

Yokouchi, Y., Ohsugi, K., Sasaki, H., & Kuroiwa, A. (1991). Chicken homeobox gene Msx-1: structure, expression in limb buds and effect of retinoic acid. *Development (Cambridge)*, *113*(2), 431-444. <https://doi.org/10.1242/dev.113.2.431>
